# Human Dicer1 hotspot mutation induces both loss and gain of miRNA function

**DOI:** 10.1101/2025.10.15.682667

**Authors:** David Jee, Seungjae Lee, Dapeng Yang, Robert Rickert, Renfu Shang, Danwei Huangfu, Eric C. Lai

**Author notes:** These authors contributed equally (David Jee, Seungjae Lee).

## Abstract

The core miRNA biogenesis enzyme Dicer1 sustains recurrent mutations in cancer that compromise its RNase IIIb domain, which cleaves 5p arms of pre-miRNA hairpins. However, the lack of knockin models has limited fuller understanding. Here, we generated *Dicer1-KO* and *Dicer1-S1344L* (homozygous and hemizygous) human ESCs; the latter is a non-catalytic mutation in RNase IIIa that impairs RNase IIIb activity. *Dicer1* knockouts lack canonical miRNAs, while S1344L induces two trends: ablation of miRNA-5p strands, and selective changes in miRNA-3p strands. Curiously, we recognized directional upregulation of miRNA-3p passenger strands, indicating a broad strand switch. We used multiple *in vitro* assays to show 3p arm-nicked pre-miRNAs preferentially load miRNA-3p species into Argonaute, compared to corresponding duplexes. Moreover, activity assays, RNA-seq data, and Argonaute-mRNA profiling, confirm that these confer increased repression capacity. These data expand the molecular consequences of *Dicer1* hotspot mutations in cancer.

## Introduction

microRNAs (miRNAs) are ∼22 nucleotide (nt) RNAs that orchestrate broad post-transcriptional regulatory networks^1^. Canonical miRNAs are generated via stepwise ribonucleolytic processing, whereby a primary miRNA (pri-miRNA) transcript is first cleaved by nuclear RNase III enzyme Drosha to release a pre-miRNA hairpin, which is then cleaved by cytoplasmic RNase III enzyme Dicer to yield a small RNA duplex. This is loaded into an Argonaute protein, which preferentially retains the guide strand (mature miRNA), while the passenger strand (miRNA* species) is discarded. Structural and sequence details of the small RNA duplex dictate selection of the mature miRNA, which can derive from either 5p or 3p arm of the pre-miRNA hairpin^2^. miRNAs guide target repression via 3’-directed mRNA decay and/or translational inhibition, via regulatory complexes recruited by the Argonaute adaptor GW182/ TNRC6.

The minimal miRNA target site consists of ∼7 nt pairing to positions 2-8 of the mature miRNA, known as the “seed” region^3,4^. Accordingly, evolutionary conservation and/or transcriptome analyses support that animal miRNAs commonly have hundreds to >1000 target sites^5^. Still, knockouts of most individual miRNA loci rarely yield overt phenotypes^6^, and the regulatory impact of predicted miRNA target sites has been challenged^7^. Detailed studies can reveal consequences in specific settings or sensitized conditions^8,9^, and ongoing work continues to reveal more about the biological impacts of miRNA mutants. However, what is not in dispute is that knockouts of core miRNA factors such as Dicer invariably leads to organismal lethality with pleiotropic defects^10–13^. In fact, *Dicer1-KO* is cell lethal in human embryonic stem cells (hESCs)^14^.

With this in mind, it is paradoxical that *Dicer1* is recurrently mutated in human cancers, implying that disabling the miRNA pathway is advantageous to tumors^15^. These “Dicer hotspot” mutations were originally recognized via heterozygous germline nonsense alleles in familial pleuropulmonary blastoma (PPB), a rare pediatric mesenchymal cancer^16^. Somatic *Dicer1* mutations were later recognized in various adult cancers, and were inevitably paired with inactivating mutations on the other allele^17^. The somatic mutations typically affect catalytic residues in the Dicer RNase IIIb domain, but not its RNase IIIa domain. As these domains cleave the 5’ and 3’ arms of the pre-miRNA hairpin^18,19^, respectively, this suggested that Dicer1 hotspot mutations selectively inactivate biogenesis of miRNA-5p species. Many studies now document specific mutation of Dicer1 RNase IIIb active sites across a range of cancers^20–25^. In addition, non-catalytic RNase IIIa mutation (S1344L) causes defects in RNase IIIb activity; this residue is physically close to the RNase IIIb active site and compromises its function^24,26^.

In previous studies, human Dicer1 variants were analyzed by plasmid transfection into surrogate mouse or human *Dicer-KO* cell lines, with *in vitro* biochemical assays, or by sequencing material from resected *Dicer1*-mutant tumors. All of these approaches have limitations for full understanding of Dicer1 hotspot mutations. These include issues related to ectopic expression, non-endogenous cell contexts, inability to assess miRNA function, contamination by wildtype cells, and challenges for normalization of genomic data.

Here, we address these issues by generating the first knock-in mutants of *Dicer1- S1344L* in human ESCs. These exhibit widespread loss of miRNA-5p species, but also an unexpected signature of selective upregulation of miRNA-3p strands. While changes in miRNA- 3p species in Dicer1 hotspot cancer were previously seen^23,24,27–30^, we elucidate that this specifically involves a maturation strand switch of passenger miRNA-3p species into functional guide miRNAs. Therefore, Dicer1 syndrome cancer is not a simple hypomorphic situation as widely conceived, but instead combines miRNA loss- and gain-of-function.

## Results

### Generation of *Dicer1* knock-out and S1344L knock-in hESC lines

In contrast to viable *Dicer-KO* mESCs^31,32^, we found that Dicer1 is essential in hESCs^14^. This necessitated a conditional strategy in which mutations were induced in a Dox-inducible *Dicer1* transgenic background (*TRE-Dicer1*). With this in mind, we sought to model *Dicer1* hotspot mutations in H1 hESCs (**Figure 1A**). We selected Dicer1-S1344L, to continue our work on how this non-catalytic RNase IIIa mutation compromises RNase IIIb catalysis^24^. We anticipated that we might recover these easily, since S1344L hemizygous tumor cells can survive and metastasize, as with conventional RNase IIIb mutant cells^24^. However, repeated CRISPR/Cas9 trials failed to recover S1344L mutant hESC lines. We therefore employed the conditional approach (**Figure 1B**), and successfully recovered both knockin and frameshift (fs) alleles, including homozygous (*S1344L*) and hemizygous (*S1344L/fs*) mutants, and null (*fs/fs*) lines. The allele sequences are provided in **Extended Data Fig. 1A**.

**Figure 1.**
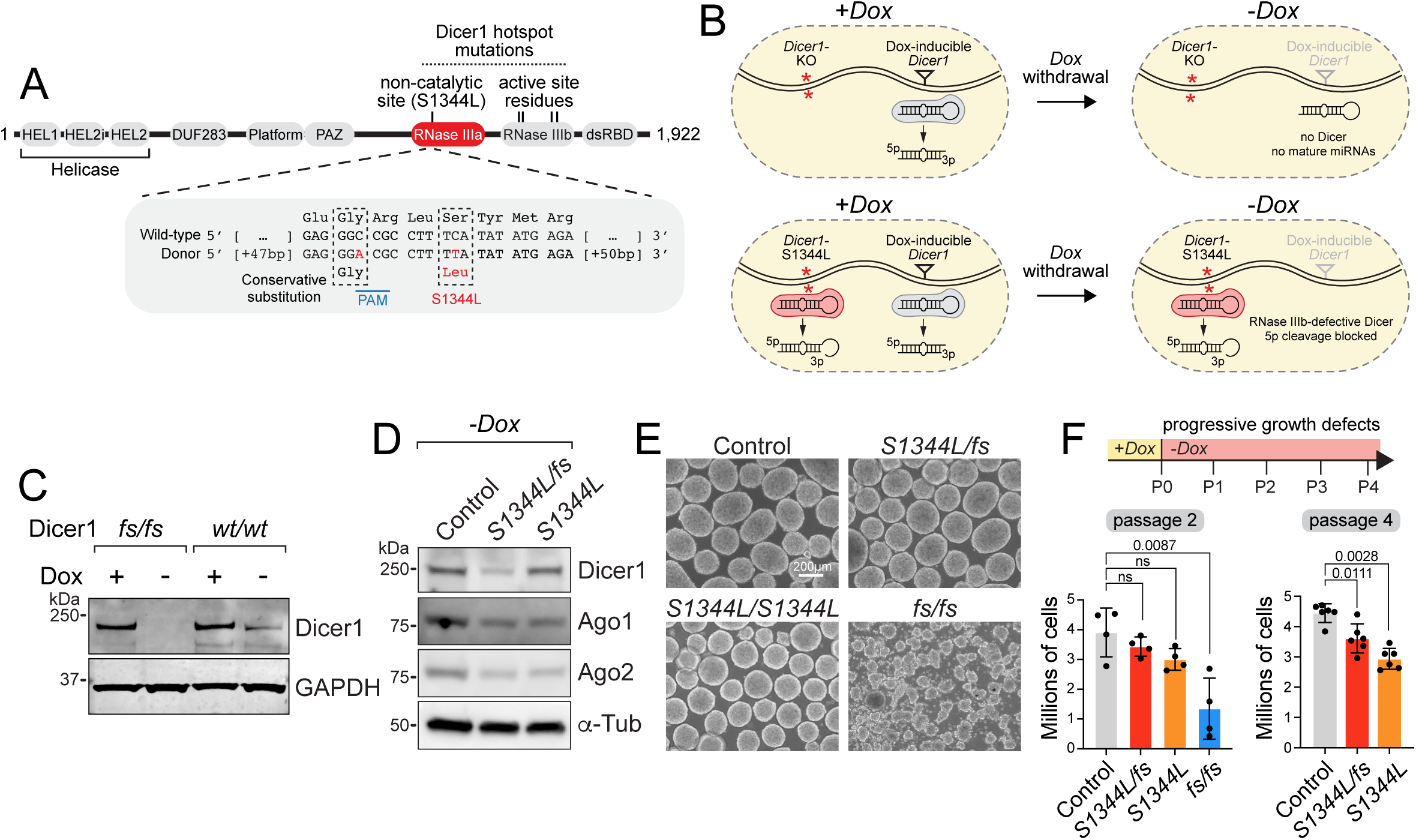
Generation and characterization of *Dicer1-KO and* S1344L human ES cells. **(A)** Domains of Dicer1, with targeted region of RNase IIIa marked. Bottom panel shows sequence of wildtype and S1344L donor. **(B)** Conditional targeting strategy. H1 hESCs express a Dox-inducible wildtype *Dicer1* transgene, allowing for successful knockin allele generation. Upon Dox withdrawal, selective activity of S1344L knockin can be studied. **(C)** Western blotting for Dicer1 shows transgene expression is under precise control of doxycycline, with complete loss of Dicer1 in *fs/fs* cells. Western blot shown was performed twice with similar results. **(D)** Western blotting in *S1344L/fs* and *S1344L* homozygous cells shows partial Dicer1 loss only in hemizygous cells, but decreased levels of Ago proteins in both S1344L cell types. Western blot shown was performed twice with similar results. Full corresponding western blot including IP lanes is shown in **Extended Data** Fig. 6A. **(E)** 3D cell culture growth under self renewing conditions shows severe growth defects in *fs/fs*, and modest defects in both *S1344L* genotypes. Scale bar, 200µm. **(F)** Progressive cell growth defects observed in *fs/fs* and *S1344L* cells. Growth curve analysis reveals severe cell growth defects in *fs/fs* cells beginning in passage 2 (left panel), with failure to survive beyond this passage. *Dicer1-S1344L* cells begin to display defects in cell growth in progressive passages beginning at passage 2, with overt differences at passage 4 (right panel). A two-tailed unpaired student’s *t*-test with Welch’s correction was applied for individual pairwise comparisons. *p*-values for each comparison are indicated. Error bars represent mean ±SD. n=4 (P2) and n=6 (P4) biological replicates.

Western blotting of parental hESCs expressing *TRE-Dicer1* showed that it moderately increased total Dicer1 protein (**Figure 1C**). By contrast, 4 days after Dox removal from *fs/fs* cells, no Dicer1 protein was detected (**Figure 1C**). Similar analysis of S1344L hESCs showed reduction of Dicer1 protein from hemizygous cells that lack one copy of *Dicer1*, but not from homozygous cells (**Figure 1D**). Thus, mutant Dicer1 accumulated normally. We showed that ongoing miRNA biogenesis is required for homeostatic stability of Argonaute proteins^33^. Both S1344L genotypes reduced Ago1 and Ago2 proteins (**Figure 1D**), suggesting that overall miRNA accumulation was compromised in these mutants.

*Dicer1*-null H1 hESCs were poorly viable in 2D culture and yielded fragmented colonies in 3D culture, phenocopying our previous HUES8 ESC mutants^14^. This was particularly noticeable in the 2nd passage following Dox withdrawal, and *Dicer1*-null H1 hESCs could not be maintained thereafter (**Figure 1E** and **Extended Data Fig. 1B**). In contrast, we could passage both types of S1344L mutant cells multiple times without Dox, and they maintained normal levels of ESC markers such as Sox2, Oct4 and Nanog (**Figure 1E** and **Extended Data Fig. 1C**). Nevertheless, S1344L mutant hESCs exhibit reduced proliferation (**Figure 1F**). The stronger effect of S1344L homozygous cells, relative to hemizygotes, hinted that total levels of mutant protein may be relevant.

As we did not recover *Dicer1-S1344L* mutants from CRISPR/Cas9 screening under competitive conditions, we infer this to reflect deleterious effects that could be overcome once stably isolated. Accordingly, the presence of conditional Dicer1 expression in our S1344L mutant lines is advantageous, as it prevents selection for compensatory mutations in these lines, and allows us to focus on proximal regulatory defects.

### Expected and unexpected defects in miRNA biogenesis in Dicer1-S1344L hESCs

Based on prior genetic and molecular studies^15,17,24^, we anticipated that mature 5p- miRNAs might be downregulated in *Dicer1-S1344L*. Still, with a point mutation in a non-catalytic residue, it was not evident how general this effect would be. We also note that prior studies of RNase IIIb mutant cancers involved sequencing of tumor RNAs followed by normalization for library size. This approach is suitable to detected biases in 5p:3p miRNA ratios. However, global distortion of miRNA levels might be masked by normalizing to a fixed read depth, and resected tumors contain variable and unknown amounts of normal cells^24^.

Our analysis of genetically characterized knockin cells, incorporating spike-in normalization, were positioned to yield quantitative data on small RNA alterations. Recognizing that miRNAs produced in the presence of active *TRE-Dicer1* may persist for days^34^, we evaluated miRNA dynamics in *Dicer1* null conditional cells using Northern blotting. Some miRNAs (e.g. miR-20a-3p) were absent within 5 days of transgene shutoff, concomitant with pre-miRNA accumulation, whereas others (e.g. miR-92a-3p) required additional days for depletion (**Extended Data Fig. 2**). We could see further depletion from 7 to 9 days, but the health of knockout cells declined at the latest timepoint. Based on this, we sequenced replicate small RNA libraries from control, knockout, and S1344L hemizygous and homozygous cells after 7 days Dox depletion.

Analysis of *Dicer1-fs/fs* hESCs showed uniform loss of both 5p and 3p miRNA populations (**Figure 2A-B, Supplementary Table 1**), indicating successful ablation of canonical miRNA biogenesis and robust normalization procedures. The rare, apparently upregulated, loci are “high-numbered” loci that do not include known Dicer-independent miRNAs (**Supplementary Table 1**). Instead, many are questionable as miRNAs, since they have non-canonical sizes, heterogeneous 5’ and/or 3’ termini, and/or explicitly overlap non-coding RNAs. Thus, *Dicer1-fs/fs* hESCs appear incompetent for canonical miRNA biogenesis, as expected.

**Figure 2.**
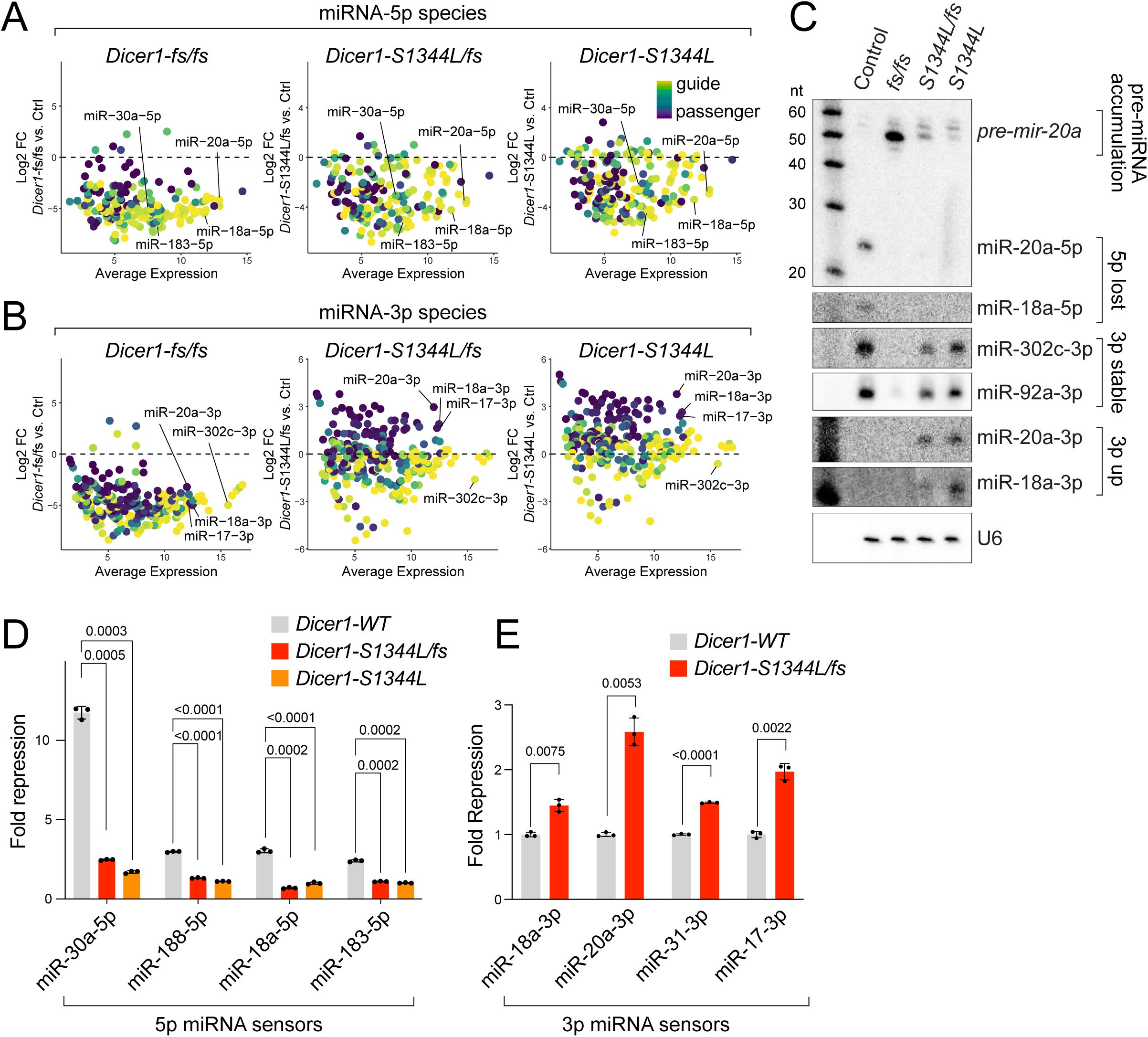
*Dicer1-S1344L* induces global miRNA-5p depletion and 3p star strand accumulation. **(A)** Spike-in normalized small RNA sequencing reveals global depletion of 5p miRNAs in *Dicer1-fs/fs*, *S1344L/fs*, and *S1344L/S1344L* cells. **(B)** Selective upregulation of passenger strand 3p miRNAs is observed in *S1344L/fs* and *S1344L/S1344L* cells. Guide and passenger strands are color coded based on proportion of 5p or 3p reads compared to total mature reads. **(C)** Northern blot validation of 5p depleted miRNAs, 3p increased miRNAs, and control unchanged miRNAs. Northern blots were repeated three times with similar results. **(D-E)** Luciferase sensor assays reveal loss of 5p miRNA activity (D) and selective upregulation of passenger 3p strand activity (E) in *S1344L* cells. A two-tailed unpaired student’s *t*-test with Welch’s correction was applied for individual pairwise comparisons. *p*-values for each comparison are indicated. Error bars represent mean ±SD. n = 3 biological replicates.

Hemizygous and homozygous S1344L cells showed similar depletion of most miRNA- 5p species (**Figure 2A**). Thus, this non-catalytic Dicer1 mutation broadly impairs RNase IIIb activity on most pre-miRNAs. The bulk of miRNA-3p species were unchanged in S1344L mutants, but we observed subpopulations that spread in either direction, including many with substantially increased levels (**Figure 2B**). We validated this using Northern blotting. *Dicer1-KO* cells exhibited uniform loss of miRNA-5p and miRNA-3p species, accompanied by increases in *pre-mir-20a* and *pre-mir-92a* (**Figure 2C** and **Extended Data** Fig. 2A). On the other hand, both S1344L hemizygous and homozygous hESCs exhibited directional loss of miRNA-5p species, while miRNA-3p species were unchanged or slightly lower in some cases, but increased with others (**Figure 2C**). Overall, compromising Dicer1 RNase IIIb activity induces complex effects on miRNA accumulation.

Previous analysis of *Dicer1* tumors showed apparent elevation in miRNA-3p species^23,24,27–30^. However in our past work, we were unable to disentangle this from skewed normalization in tumor sequencing, and we did not validate strand switching for miRNA-3p species tested^24^. Nevertheless, an aspect of this trend proved true, and following studies clarify both its scope and underlying mechanism.

### Dicer1-S1344L elevates the expression and activity of miRNA-3p passenger species

*Dicer1* mutant cells presumably incur a range of primary and secondary effects. For example, RNA-seq analysis of Dicer1 hotspot endometrial cancers revealed target upregulation for certain miRNA-5p and miRNA-3p species^24^. While this may partly be due to transcriptional changes at miRNA genes, this would not explain the broad accumulation of miRNA-3p species in Dicer1-S1344L hESCs (**Figure 2B**). Upon inspecting these loci, we realized our Northern validations involved strand switches (**Figure 2C**). That is, with *mir-18a* and *mir-20a*, their respective miRNA-5p species accumulated in wildtype, but their miRNA-3p species accumulated in *Dicer1* mutants. This indicated loss of mature miRNA species and misexpression of miRNA* species (i.e. passenger strands), including ones that do not accumulate in wildtype.

We tested if this reflected a broader trend in RNase IIIb-deficient hESCs. To do so, we color-coded the expression of miRNA-5p and miRNA-3p species according to their strand bias in wildtype (5p/total reads). This made it visually apparent that miRNA-5p species were downregulated in mutant hESCs, regardless of their functional status, but that miRNA-3p species were directionally elevated if they were miRNA* species (**Figure 2A-B**). There was also moderately biased downregulation of some miRNA-3p guide strands, in accord with Northern blotting (**Figure 2C**). We found these trends in both homozygous and hemizygous *Dicer1- S1344L* hESCs, shown in MA plots (**Figure 2A-B**) and volcano plots (**Extended Data** Fig. 3). We also assessed specificity of terminal cleavage sites of 5p and 3p small RNAs, but did not observe substantial differences (**Extended Data** Fig. 4). Thus, Dicer1 hotspot mutants primarily affect small RNA expression, but not the precision of Dicer-defined ends.

Accumulation of small RNAs does not necessarily imply elevated activity. For example, we showed that accumulation of the star species miR-486-3p in Ago2-catalytic dead cells actually reflects trapping of non-functional duplex intermediates with mature miR-486-5p^35^. We occasionally detected endogenous species corresponding to miRNA-5p+loop sequences in the presence of Dicer1-S1344L (e.g. *mir-92a*, **Extended Data** Fig. 2B); however, we did not reliably detect these for other loci (**Figure 2C**). Others also observed these, primarily using *in vitro* Dicer processing reactions^20,21,36–38^, but they seem difficult to detect without overexpression of mutant Dicer and/or miRNA misexpression^18,24^. We take this to reflect their preferential degradation, as opposed to residing within stable nicked hairpins with their companion miRNA-3p species.

These implied that miRNA-3p passenger strands might truly be increased as mature species. We conducted luciferase sensor assays to probe changes in miRNA activities. Most miRNA-5p species exhibited strongly decreased repression capacity in Dicer1-S1344L hESCs (**Figure 2D**), consistent with their expression changes. Reciprocally, we validated increased activity of multiple miRNA-3p species (**Figure 2E**). Several of these (miR-17a-3p, miR-18a-3p, miR-20a-3p) derive from the “oncomir” cluster (*mir-17∼92*), which is well-expressed in hESCs, but we also detected increased activity of non-cluster loci (miR-31-3p). Thus, inactivation of Dicer1 RNase IIIb function not only broadly depletes miRNA-5p species, it causes a non-intuitive increase in functional miRNA* species derived from 3p hairpin arms. These findings unify and explain prior observations on deregulated miRNA-3p expression in Dicer1 hotspot _cancer_23,24,27-30_._

### 3p-nicked pre-miRNA hairpins exhibit enhanced maturation of miRNA-3p RISC

We evaluated how loss of Dicer1 RNase IIIb activity might switch strand selection. We applied three complementary *in vitro* assays using radiolabeled substrates to interrogate maturation of the RNA induced silencing complex (RISC). These include (1) analysis of RISC assembly on native PAGE, (2) immunoprecipitation of assembled Ago2 complexes followed by native PAGE, and (3) unwinding assay of RISC complexes (**Extended Data** Fig. 5A). Such assays were used to determine features that control guide strand selection^39^, including 5’ nucleotide identity (favoring 5’-U) and terminal duplex asymmetry (favoring the more thermodynamically unstable end)^40,41^. Notably, *in vitro* assays can dissociate indirect consequences on small RNA accumulation that might occur in cells.

We inferred that loss of mature miRNA-5p species in Dicer1-S1344L hESCs was accompanied by transient presence of miRNA-5p+loop species produced from pre-miRNA nicking on the 3p arm by RNase IIIa activity. Although these do not accumulate stably, we detected them from endogenous (**Extended Data Fig. 2**) and ectopically expressed^24^ miRNAs in RNase IIIb mutant conditions, and they were detected from *in vitro* dicing reactions^20,21,36–38^. Accordingly, we tested whether guide strand selection was altered when utilizing pre-miRNA hairpins nicked on the 3p arm at the RNase IIIa cleavage site, compared to conventional Dicer-cleaved small RNA duplexes (**Figure 3A**). We tested miRNA-5p loci that exhibit robust strand switching in the presence of Dicer1-S1344L (*mir-20a* and *mir-30c-2*), and control miRNA-3p loci that do not alter strand selection in mutants (*mir-95* and *mir-25*) (**Figure 3B**). The positions of pre-miRNA nicks are designated in these schematics (**Figure 3B**).

**Figure 3.**
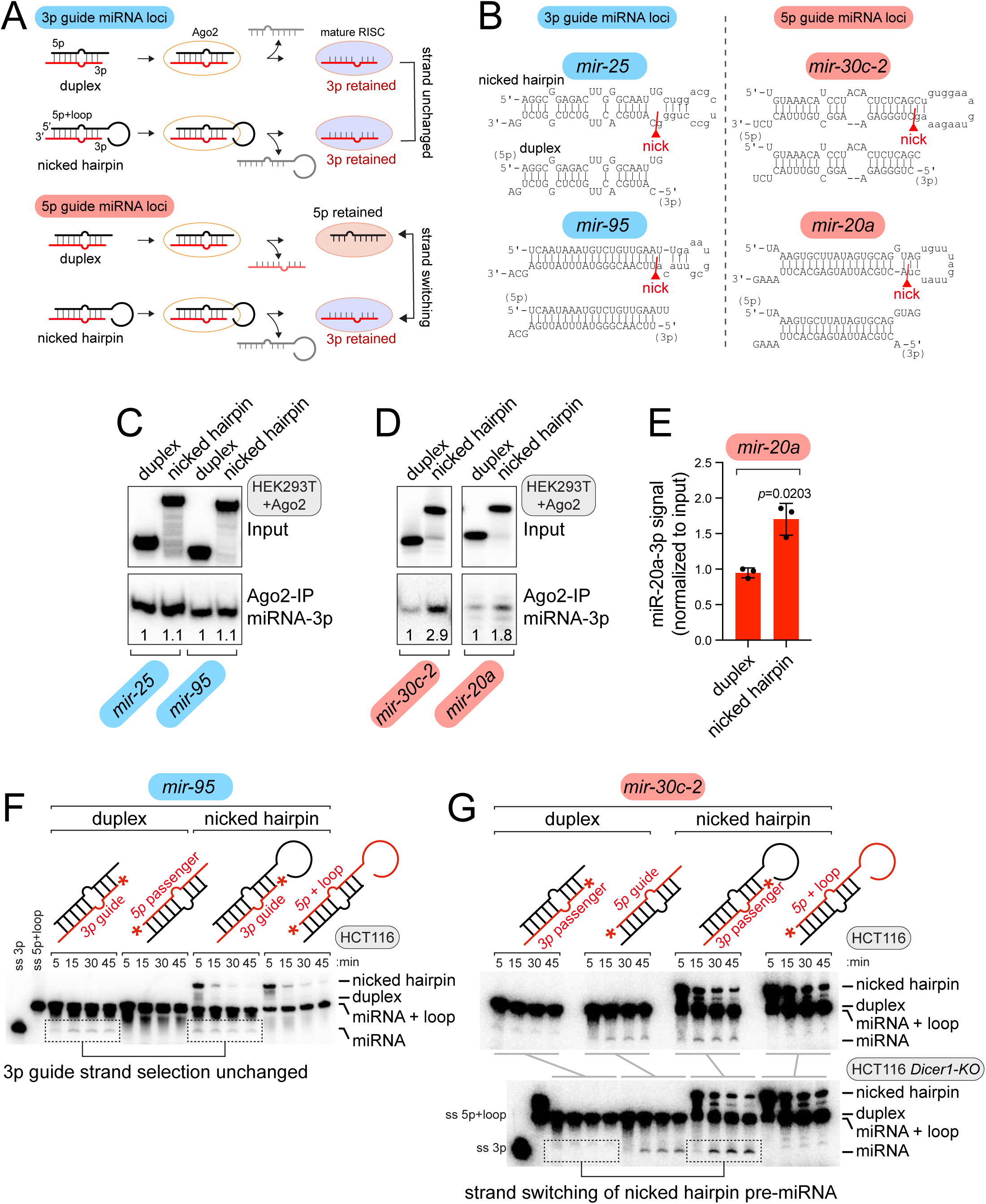
3p-arm nicked pre-miRNAs preferentially load passenger 3p strands into Ago2. **(A)** Schematic of outcomes with *in vitro* RISC assays, comparing behavior of 3p guide vs. 3p passenger strands, between duplex and nicked hairpin substrates. **(B)** Secondary structures of miRNA-3p guide (highlighted in blue throughout) and miRNA-5p guide (highlighted in red throughout) substrates tested in *in vitro* RISC assays. **(C-E)** Ago2 loading assays using Ago2- overexpressing HEK293T lysates. Substrates were incubated with lysate at 30°C for 30 min, followed by Ago2-IP. Associated RNAs were purified and analyzed on 15% native PAGE gels. **(C)** Control Ago2 loading experiments using 3p guide miRNA substrates. Similar 3p incorporation into Ago2 complexes is obtained using duplex and nicked hairpin substrates. Assay shown was repeated twice with similar results. **(D)** Ago2 loading assays with 5p guide (3p-passenger) miRNA loci shows strand switching between duplex substrate (5p guide selected) and nicked hairpin substrates (3p guide selected). **(E)** Quantification of Ago2-loaded 3p products across 3 independent assays of *mir-20a*. Significant difference between the two substrates were assessed using a two-tailed unpaired *t*-test with Welch’s correction, and the *p*- value is indicated. Error bars represent the mean ±SD. **(F-G)** Unwinding assays using wildtype and/or *Dicer1-KO* HCT116 lysates, and all four possible 5p and 3p labeled duplex and nicked pre-miRNA substrates. Reaction products were analyzed on 15% SDS-polyacrylamide gels. Assay shown was repeated three times with similar results. **(F)** Control unwinding assays using a 3p-guide locus. miR-95-3p is preferred over miR-95-5p as the guide species, from both duplex and nicked hairpin substrates. **(G)** Unwinding assays using a 5p-guide locus shows switching of 5p maturation from a duplex to preferred 3p maturation from the equivalent nicked pre-miRNA hairpin. Strand switching of 3p passenger species is independent of Dicer1 (lower panel).

We previously performed RISC assembly assays by incubating radiolabeled substrates with wildtype HEK293T cell lysates^42^, which we separated on 4% native gels (**Extended Data Fig. 5B**). We performed preliminary assays using wildtype and *Dicer1-KO* HCT116 cell lysates to compare our previously utilized substrate *Pp-luc* duplex with *mir-95* duplex. Both substrates yielded a similarly sized major product, whose intensity was stronger when using *Dicer1-KO* lysates (**Extended Data Fig. 5C**). This identified mature “holo” RISC, which assembles more efficiently *in vitro* when endogenous miRNA biogenesis is blocked. We also note that presumed intermediate complexes exhibited different sizes with different substrates (**Extended Data Fig. 5C**). We infer these are pre-RISC, but as we are not confident of their composition, we do not specifically label them in the figures.

To focus on intrinsic capacity of Argonaute loading, we utilized lysates from *Dicer1-KO* HEK293T cells^43^ that we transfected with Ago2 plasmids. As these assays utilize the same 5’- radiolabeled miRNA-3p species in duplex and nicked hairpin configurations, we can directly compare effects of precursor structure on RISC assembly. With duplex loading, relatively little RISC formed in this timecourse. However, parallel experiments using a labeled 3p-guide within a nicked hairpin substrate exhibited enhanced maturation (**Extended Data Fig. 5D-E**). Thus, hairpins can be valid substrates for RISC assembly, as documented with canonical pre-miRNAs^44^ and as is obligately the case for Dicer-independent *mir-451*^45–47^.

We compared this behavior to miRNA-3p species that are normally passenger strands, but accumulate robustly in small RNA sequencing and/or Northern blotting (i.e., miR-20a-3p and miR-30c-2-3p). Notably, when the miR-20a-3p arm was labeled, it efficiently matured as the guide strand from the nicked hairpin, compared to the duplex substrate (**Extended Data Fig. 5D-E**). We observed similar results using *mir-30c-2*. These *in vitro* tests provide direct evidence for strand switching in RISC assembly from unconventional 3p-nicked hairpins.

A limitation of RISC assembly assays is that they do not specifically reflect incorporation into Ago2. To do so, we transfected myc-tagged Ago2 into HEK293T cells and prepared lysates for RISC assembly, from which we immunoprecipitated (IP-ed) Ago2, isolated its associated RNAs, and visualized them on 15% native PAGE. We established this approach by testing 3p guide miRNA substrates (**Figure 3B**). Both miR-25-3p and miR-95-3p were efficiently IP-ed with Ago2, in a manner that was similar between duplex and nicked hairpin substrates (**Figure 3C**). However, the behavior of normally passenger strand species miR-30c-2-3p and miR-20a-3p was different. In both cases, a greater proportion of input was incorporated into Ago2 when using the nicked hairpin substrate, compared to conventional duplexes (**Figure 3D**). We quantified the increase of mature miR-20a-3p in Ago2 complexes from nicked hairpins from three trials (**Figure 3E**).

Finally, we performed duplex unwinding to monitor conversion of the total in vitro RNA pool into mature RNA products. This utilizes the same reaction conditions to assemble RISC complexes, but resolves reaction products on SDS-PAGE to denature proteins, but not RNA species. Here, we compared the properties of 5p and 3p radiolabeled substrates. With *mir-95*, a control 3p guide substrate, we observed preferred retention of single-stranded miR-95-3p from both duplex and nicked substrates (**Figure 3F**). However, with *mir-30c-2*, normally a 5p guide substrate, we observed opposite behavior of these precursors. The radiolabeled 5p mature strand was produced from the duplex but not the nicked hairpin, while the radiolabeled 3p mature strand was generated from nicked hairpin but not duplex precursor (**Figure 3G**). We tested the panel of *mir-30c-2* substrates using *Dicer1-KO* lysates, and obtained similar strand-switching results (**Figure 3G**).

Altogether, the three *in vitro* assays support aberrant 3p passenger strand maturation from nicked hairpins present in *Dicer1* hotspot mutant cells, but this does not depend on Dicer1 *per se* and appears to be an intrinsic property of Argonaute.

### *Dicer1-S1344L* hESCs exhibit global alterations in Ago2 targeting

As we demonstrated both gain and loss of miRNAs in Dicer hotspot conditions, we characterized effects on the transcriptome. This was relevant as miRNA abundance in small RNA sequencing may not necessarily reflect active RISC. Therefore, we prepared enhanced crosslinking immunoprecipitation (CLIP) libraries for Ago2, from 11 day Dox-depleted wildtype, S1344L *pm/fs* and *pm/pm* cells in biological replicates.

We tested immunoprecipitation (IP) of crosslinked samples using pan-AGO antibody and Ago2-specific antibody. While western blotting showed that pan-AGO antibody pulled down AGO complexes, it was less efficient than Ago2 antibody based on autoradiography of crosslinked, fragmented, and labeled RNA (**Extended Data Fig. 6**). Thus, we cloned materials from Ago2-specific pulldown. While Ago2-IP recovered less Ago2 protein and less crosslinked RNA from S1344L mutants than wildtype (**Extended Data Fig. 6**), we were able to clone biologically replicate CLIP libraries for each genotype. A caveat is that due to lower RNA recovery from *Dicer1* mutants, more amplification was required to recover libraries, which might introduce some bias (see **Methods**). Accordingly, we performed several quality checks.

As mature miRNA reads could be quantified from Ago2-CLIP libraries, we first compared 5p and 3p miRNA changes. miRNA expression in total small RNA and Ago2-CLIP libraries was broadly concordant (**Extended Data Fig. 7A**). In particular, Ago2-CLIP data reproduced systematic depletion of miRNA-5p species and selective increases in miRNA-3p passenger strands, in both hemizygous and homozygous S1344L mutants (**Figure 4A-B, Supplementary Table 2**).

**Figure 4.**
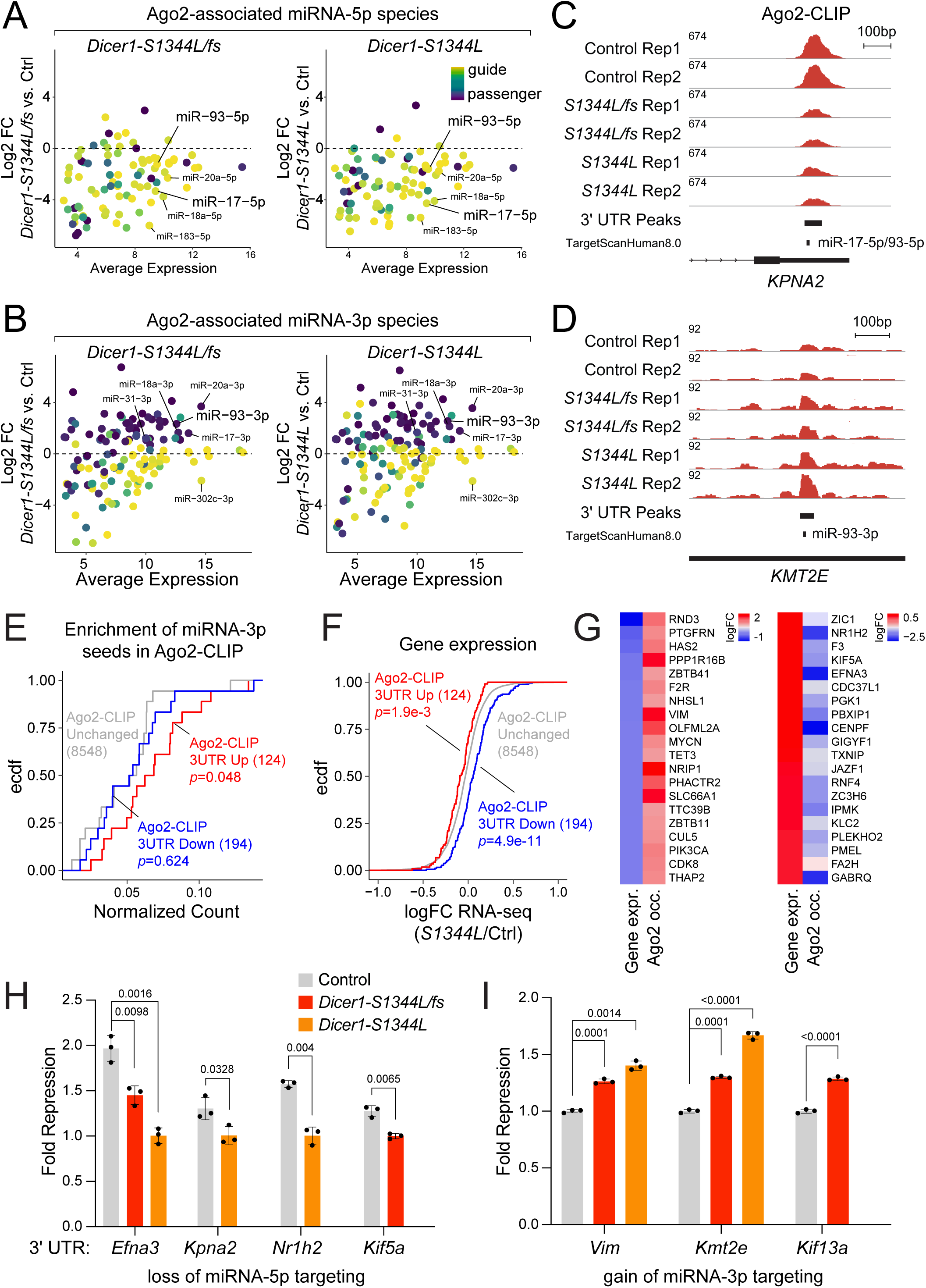
Ago2-CLIP reveals decreased targeting by 5p-miRNAs and ectopic targeting by 3p- miRNA passengers. **(A-B)** Ago2-associated miRNAs from Ago2-CLIP data reveal **(A)** directional 5p-miRNA loss and **(B)** selective increase of passenger strand 3p-miRNAs, consistent with total small RNA-seq. 5p and 3p miRNAs are color coded based on proportion of respective reads relative to total mature reads. **(C)** IGV browser screenshot of *KPNA2* 3’ UTR showing depletion of CLIP occupancy at miR-17/93-5p seed target site. **(D)** 3’ UTR of *KMT2E* shows increased Ago2 CLIP signal centered on miR-93-3p seed site, consistent with associated change in miRNA level. **(E)** CDF plot of distribution of genes based on frequency of seed sites in 3’ UTRs associated with increased 3p-miRNAs shows enrichment in 3’ UTRs harboring increased Ago2 targeting. Detected seed numbers were normalized by the corresponding 3’ UTR length. Kolmogorov- Smirnov test was used to determine *p*-value of separation between control and test sets. **(F)** CDF plot of expression changes from RNA-seq in genes harboring increase or decrease in total Ago2-CLIP occupancy on 3’ UTRs. These are associated with overall decreased or increased gene expression in *Dicer1-S1344L* mutant cells. Kolmogorov-Smirnov test was used to determine *p*-value of separation between control and test sets. **(G)** The top 20 genes that are reciprocally regulated by miRNAs, identified by changes in Ago2-CLIP 3’ UTR occupancy in S1344L mutant cells. **(H-I)** Experimental validation of altered miRNA functions in S1344L mutant cells. 3’ UTR luciferase sensors were constructed for genes with reduced miRNA-5p targeting **(H)** and increased miRNA-3p targeting **(I)** from Ago2-CLIP data. A two-tailed unpaired student’s *t*-test was applied for individual pairwise comparisons. *p*-values for each comparison are indicated. Error bars represent mean ±SD. n=3 biological replicates.

We next assessed CLIP library coverage across coding sequences (CDS), 3’ untranslated regions (3’ UTRs), 5’ UTRs, and intergenic regions. Ago2 was enriched in 3’ UTRs, consistent with the dominant targeting sites of miRNAs (**Extended Data Fig. 7C-D**). We then analyzed Ago2-CLIP coverage across predicted conserved seed matches to expressed hESC miRNAs. A number of miRNAs exhibit focal Ago2-CLIP peaks centered on predicted sites, and we observed cases in which downregulated miRNA-5p species in S1344L mutant cells were associated with decreased Ago2-CLIP signals. However, we observed counter-examples, e.g. that Ago2-CLIP occupancy was not focally enriched at seed matches. There are several possible explanations for this. The efficacy of miRNA targeting inferred from transcriptome studies frequently does not align with Ago2-CLIP occupancy^48^, possibly because cross-linking freezes Ago2 on transient sites. Reciprocally, computationally predicted sites may not be stably bound by Ago2 in cells. Nevertheless, given enrichment of Ago2-CLIP signals on 3’ UTRs, and appropriate behavior of candidate target sites, we assessed these data further.

We observed sets of miRNA-5p targets with decreased Ago2 occupancy (**Figure 4C, Extended Data** Fig. 8**, Supplementary Table 2**), as expected from the small RNA sequencing and Ago2-associated miRNAs. We note that decreased miRNA-5p expression was not always mirrored by decreased Ago2 association in S1344L mutants. It was previously shown that genetic knockouts of miRNAs did not necessarily eliminate Ago2 binding to cognate sites, possibly due to association via other miRNAs^49^. Still, these observations were consistent with compromised miRNA-5p biogenesis in S1344L mutants (**Figure 2**).

Of greater interest was to assess targeting via aberrant miRNA-3p species in Ago2 complexes. Although passenger strands harbor detectable regulatory activity in mammalian cells^50^, target predictions for passenger strands are not robust, and thus are omitted from most prediction tools. Still, we could identify genes with concomitant increase in Ago2-CLIP occupancy at seed matches to increased miRNA-3p species in S1344L mutants (**Figure 4D** **and Extended Data** Fig. 8B).

Such Ago2 peak changes were most apparent when examining isolated peaks, but many genes exhibit complex alterations of Ago2-CLIP signals, with CLIP regions that frequently overlap multiple predicted seed matches. This motivated us to calculate aggregate Ago2-CLIP signals across 3’ UTRs (**Extended Data** Fig. 7**, Supplementary Table 5**), allowing us to subset those genes with overall decreased or increased Ago2 occupancy. This is rational, as miRNAs of different seeds will collectively contribute to repression of an individual transcript. This proved informative when examining the distribution of miRNA-3p seed matches in genes with aggregate alteration of Ago2-CLIP occupancy in S1344L mutants. Genes with net decreased Ago2-CLIP 3’ UTR signals in mutants showed similar distribution of miRNA-3p seed matches, but genes with net increased Ago2-CLIP 3’ UTR signals exhibited higher frequency of miRNA- 3p seed matches (**Figure 4E**).

Encouraged by these trends, we generated RNA-seq data (**Extended Data** Fig. 7A**, Supplementary Table 4**), and integrated this with Ago2-CLIP data. Here, we focused on *S1344L* homozygous cells, to avoid the variable of *Dicer1* heterozygosity in *S1344L/fs* cells. Interestingly, loci with net decrease in 3’ UTR Ago2-CLIP occupancy shifted towards higher expression, while genes with net increased 3’ UTR Ago2-CLIP signals shifted towards lower expression (**Figure 4F**). Examples of genes with reciprocal trends in Ago2-CLIP and gene expression in *S1344L* mutants are shown in **Figure 4G**. These data support the notion that a subset of genes in Dicer-S1344L have enhanced repression due to elevated Ago2 association, via increased miRNA-3p species.

We selected targets for experimental validation. For this purpose, we focused on genes with individual, focal, Ago2-CLIP signals at seed matches, and generated a panel of luciferase 3’ UTR sensors (**Figure 4H-I** and **Extended Data** Fig. 9 and **Supplementary Table 3**). These include genes with loss of miRNA-5p targeting (*Efna3*, *Kpna2, Nr1h2*, *Kif5a*) and ones with increased miRNA-3p targeting (*Vim*, *Kmt2e*, *Kif13a*) in S1344L mutants.

To test these, we cultured wildtype and S1344L hESCs without doxycycline for 6 days, before transfecting each sensor. We cultured cells for two more days without Dox, before assessing luciferase activities. As 3’ UTR sensors are often only mildly repressed by miRNAs, we took care to compare their response with corresponding cells in the presence of doxycycline. Under these conditions, expression of the *Dicer1* transgene is expected to normalize effects of the endogenous *Dicer1* mutations. Although hESCs are modestly transfected using lipofectamine, we obtained reproducible data with these sensors.

In particular, 3’ UTR sensors for genes with reduced association of Ago2-CLIP at miRNA-5p seed matches exhibited reduced repression in S1344L mutants (**Figure 3H**). Reciprocally, 3’ UTR sensors for genes harboring increased Ago2-CLIP signals at discrete miRNA-3p seed matches were subject to enhanced repression in S1344L mutants. Although these trends were modest, we emphasize these sensors respond to endogenous hESC miRNAs. We further validated responses of these 3’ UTR sensors to cognate miRNAs, as standard luciferase assays with ectopically expressed miRNAs in HEK293T cells showed dose-sensitive repression (**Extended Data Fig. 9**).

Overall, these computational and experimental assays demonstrate that compromising Dicer1 RNase IIIb function results not only in loss-of-function of miRNA-5p species, but also gain-of-function of miRNA-3p passenger strands.

### *Dicer1*-hotspot tumors exhibit opposite effects on targets of miRNA-5p and -3p species

We investigated signatures of altered miRNA-mediated regulation in tumors. Our previous survey of TCGA data^51^ showed that the uterine corpus endometrial carcinoma (UCEC) cohort harbored numerous RNase IIIb mutant samples^24^. We used 15 Dicer1 hotspot and ∼490 control UCEC cases for differential gene expression and miRNA expression analysis. We anticipated it might be challenging to observe opposite effects of miRNA-5p and -3p species on gene expression. First, we showed that only a few miRNA seeds showed directional upregulation of predicted targets, and while some were miRNA-5p species, others were miRNA- 3p species^24^. This suggested there were indirect effects across tumor transcriptomes, which is inevitable. Second, we inferred that signals of upregulated miRNA-3p species in UCEC tumors are in part likely a normalization artifact^24^, due to heterogeneity of tumor samples and lack of spike-ins in TCGA small RNA data^52^. Third, we did not have Ago2-CLIP data to stratify these gene sets in tumors, and therefore had to rely fully on target predictions.

Nevertheless, given our data on passenger strand switching, we attempted to derive signals for altered targeting by 5p and 3p miRNAs. To this end, we compared gene sets as defined by the frequency of seed matches to 5p or 3p miRNAs, whose expression levels were altered between Dicer1 hotspot and control tumors in the TCGA UCEC cohort. When testing downregulated 5p miRNAs in hotspot tumors (log_2_FC < -1) (**Figure 5A** and **Supplementary Table 6**), we observed robust cumulative derepression that correlated with numbers of 5p seeds (**Figure 5B**) and strength of seed types (8mer > 7mer-m8 > 7mer-A1) (**Extended Data Fig. 10A**).

**Figure 5.**
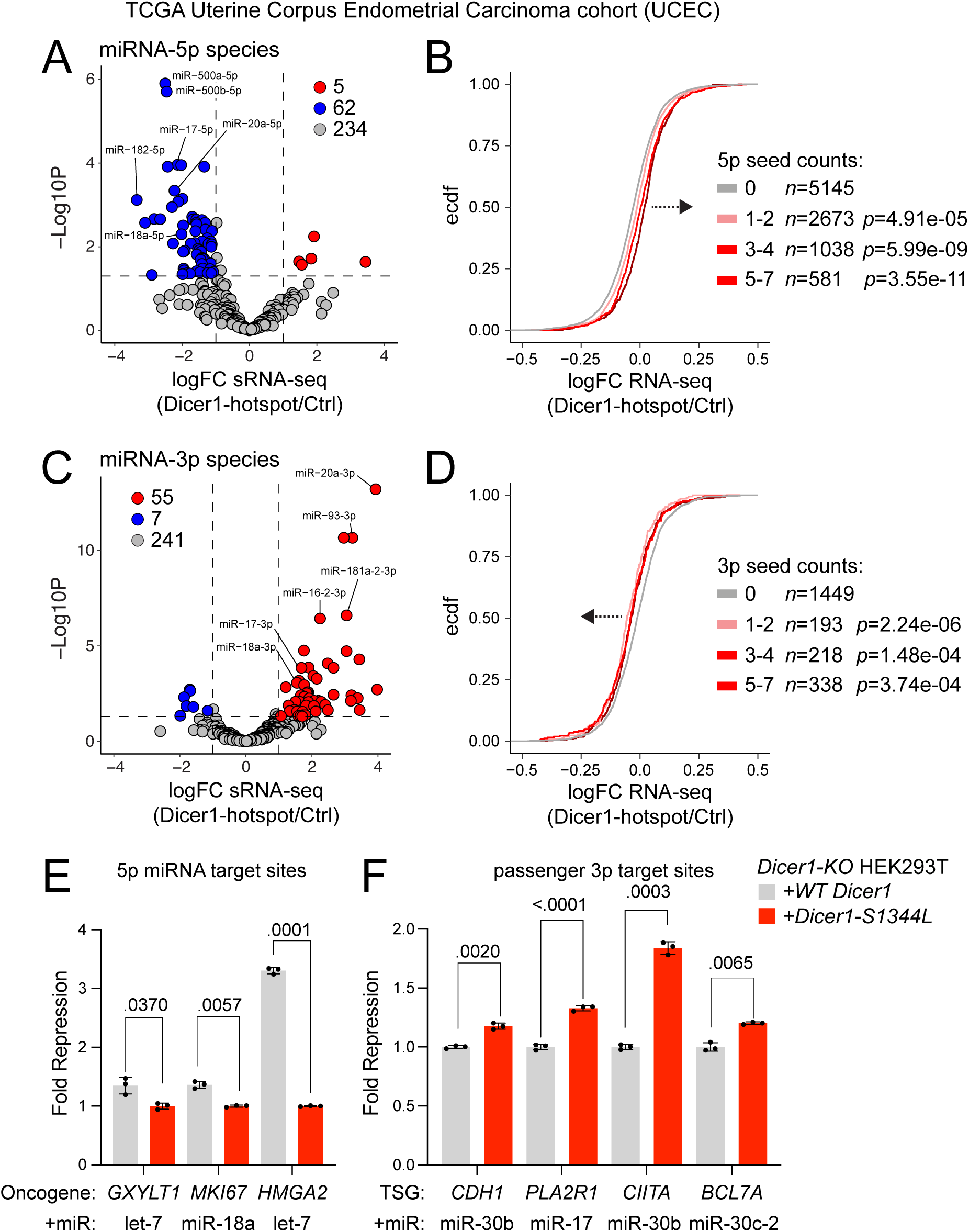
Gene expression in TCGA Uterine Corpus Endometrial Carcinoma Dicer1 hotspot tumors. **(A)** Comparison of 5p miRNA levels in Dicer1 hotspot and control TCGA UCEC cases reveal global depletion in hotspot tumors. **(B)** Derepression of miRNA-5p targets in UCEC RNA-seq data. Genes were grouped into control set (gray line with 0 count of 5p seed matches) and test sets (pink series with increasing counts of 5p seeds in 3’ UTRs). Numbers of genes in each category are shown along with boxed gene set categories. Kolmogorov-Smirnov test was used to determine *p*-value of separation between control and test sets. **(C)** Comparison of 3p miRNA levels in Dicer1 hotspot and control TCGA UCEC cases reveal selective increase in hotspots. **(D)** Genes were grouped into control set (gray line with 0 count of 3p seeds) and test sets (pink series with increasing counts of 3p seed matches). miRNA-3p targets are downregulated in Dicer1 hotpot tumors. Kolmogorov-Smirnov test was used to determine *p*-value of separation between control and test sets. (**E-F**) Luciferase 3’ UTR sensor assays of cancer-relevant target genes, using Dicer1-KO HEK293T cells that were co-transfected with wt (gray) or S1344L (orange) Dicer1 plasmids and the indicated miRNA expression constructs. (**E**) Sensor assays of oncogene 3’ UTRs with decreased miRNA-5p species. (**F**) Sensor assays of tumor suppressor 3’ UTRs with increased miRNA-3p species. These tests validate opposite effects of 5p guide and 3p passenger strands in Dicer1 hotspot conditions. A two-tailed unpaired student’s *t*-test with Welch’s correction was applied for individual pairwise comparisons. *p*-values for each comparison are indicated. Error bars represent mean ±SD. n=3 biological replicates.

We next addressed potential impacts of 3p seed frequency to differential gene expression in UCEC Dicer1 hotspot versus control tumors. Although passenger strands often have highly conserved sequences and can mediate repression^53^, even in mammals^50^, their effects on gene regulation are overall modest compared to guide strand miRNAs. Nevertheless, we detected an opposite signature as seen with depleted 5p miRNAs, with targets of increased 3p passenger miRNAs (**Figure 5C** and **Supplementary Table 6**) exhibiting cumulative shifts for increased repression, compared to control genes lacking any 3p sites (**Figure 5D**). We detected such signal across a range of increasing numbers of 3p sites (up to 30 occurrences in given 3’ UTRs). However, further increases in 3p site abundance led to a non-intuitive loss of repression signal (**Extended Data Fig. 10B-D)**. One might expect to observe greater repression with greater numbers of target sites. However, we realized this might be explained if there were concomitant increases in co-occurring 5p sites on highly targeted 3’ UTRs. Indeed, genes with low numbers of predicted 3p sites tended to have few 5p sites, but a multiplicity of 5p sites was frequent in genes that we categorized with large numbers of 3p passenger strand matches (**Extended Data Fig. 10D**). This rationalizes the dilution of 3p regulatory effects with large numbers of sites.

We exploited this observation to extract 3p activity signature in gene sets with high numbers of 3p passenger sites. By excluding genes with 5p seed matches, we could partially recover enhanced repression of genes with large number of 3p sites (**Extended Data** Fig. 10D). This indicated that signals for over-active miRNA-3p species may be diluted by confounding effects of broad miRNA-5p loss in Dicer1 hotspot tumors.

### Evidence for opposite effects on cancer-relevant targets of miRNA-5p and -3p species in Dicer1 hotspot settings

We began this study by noting that a *Dicer1* hotspot mutation is deleterious in self-renewing hESCs (**Figure 1**), and not overtly tumor suppressive. Therefore, we sought evidence for opposite effects on accumulation and activity of miRNA-5p species and miRNA-3p passenger strands on cancer-relevant targets. To do so, we queried the UCEC cohort for targets that (1) increase in Dicer1 hotspot cases, bear multiple sites for depleted 5p miRNA(s), and constitute oncogenes as defined by COSMIC^54^, or (2) decrease in hotspot cases, bear multiple sites for increased 3p miRNAs, and are classified as tumor suppressors. In this manner, we selected 3 oncogenes and 4 tumor suppressors for tests.

Since an accepted cell model for UCEC is not available, and hESCs are relatively poorly transfected, we used HEK293T cells as a test vessel. We previously showed that *Dicer1-KO* HEK293T cells^43^ can be transfected with wildtype or S1344L Dicer plasmids to recapitulate selective defects in miRNA biogenesis^24^. Importantly, while our prior tests of *mir-151* and *mir- 199a-1* showed loss of miRNA-5p species but no effects on miRNA-3p accumulation, both of these yield mature miRNA-3p species, which are not expected to change in light in of our current data. To sensitize these assays, we included expression constructs for the miRNA with the most sites on these 3’ UTRs (**Extended Data Fig. 10E**).

These tests revealed oppositely directed patterns of target repression between the two groups. That is, multiple oncogene 3’ UTR sensors were upregulated in the presence of S1344L vs. wildtype Dicer1 (**Figure 5E**), while tumor suppressor 3’ UTR sensors were downregulated in the mutant condition (**Figure 5F**). Although the quantitative effects were modest for some 3’ UTR sensors, the effects were significant and consistent in direction. While it is clearly simplistic to suppose that all oncogenes are up and all tumor suppressors are down in *Dicer1* hotspot cancer cells, our data support that the reciprocal consequences of mutant Dicer1 on miRNA-5p and passenger-3p species could have dual regulatory effects in tumors.

## Discussion

While specific alterations of Dicer1 activity are selected for in cancer, their consequences on miRNA biogenesis were not fully understood. Here, we use mutant hESC models to elucidate how a point mutation that inactivates Dicer RNase IIIb has complex consequences on miRNAs. This includes global loss of miRNA-5p function, as well as broad strand-switching of miRNA-3p passenger strands into functional guide strands that impact gene regulation. Therefore, Dicer1 hotspot cancer settings are not simply selective hypomorphs for miRNA function, as currently envisioned, but instead harbor neomorphic miRNA activity. This knowledge is relevant to interpret molecular consequences in Dicer1 pathologies.

The full appreciation of these biogenesis defects depended on precise knock-in models and rigorous spike-in normalized small RNA library analysis. While we and others have hinted at potential miRNA-3p upregulation in Dicer1 RNase IIIb hotspot conditions^23,24,27–30^, in our previous studies, we could not attribute our finding of miRNA-3p upregulation to functional alteration in Dicer1, as opposed to normalization artifacts in the absence of spike-in references, tumor heterogeneity^24^. Here, we clarify specific upregulation of miRNA-3p species that were normally passenger strands, and link this to ectopic target gene suppression.

Importantly, we find that loading of miRNA hairpins nicked on the 3p arm are efficiently associated with RISC. Although aberrant, such intermediates should be the dominant state of miRNA hairpins in Dicer1 hotspot cancers. From a mechanistic viewpoint, the broad strand switching in Dicer1 RNase IIIb mutant settings is unique. While 3’ untemplated uridylation can switch the functional miRNA product of *mir-324*^55^, and heterogeneous strand-switching has been noted in cancer^56^, this is the first large-scale, directional, switch in strand preference observed in a specific human mutant.

Our conditional *Dicer1-S1344L* human ESCs complement recent *Dicer* RNase IIIb in mice^30^, comprising valuable genetic tools. In particular, our knock-in hESC models for Dicer1 syndrome enable modeling of cell states and identities relevant to Dicer1 pathologies. A notable finding is that inactivation of Dicer1 RNase IIIb is overtly deleterious in hESCs. Thus, those rare cancers that enrich for Dicer1 RNase IIIb mutations, including pleuropulmonary blastoma, thyroid neoplasia, sarcomas, and uterine cancers^15,24,57^, may express specific miRNAs whose modulation contributes to tumor progression. In the future, these hESC models can be differentiated along different lineages to evaluate settings where loss Dicer1 RNase IIIb confers cell advantages.

## Methods

### Cell Line and Maintenance

All experiments involving hESCs were approved by the Tri-SCI Embryonic Stem Cell Research Oversight Committee (ESCRO) and regularly confirmed to be mycoplasma-free by the MSKCC Antibody & Bioresource Core Facility. hESCs were maintained in Essential 8 (E8) medium (Thermo Fisher Scientific, A1517001) at 37°C with 5% CO_2_ and doxycycline (2 μg/mL) were kept continuously. For the 2D culture, hESCs were seeded with 250,000 cells/well on a vitronectin (VTN, Thermo Fisher Scientific, A14700) pre-coated 6-well plate and passaged every 3∼4 days using 0.5 mM EDTA (KD Medical, #RGE-3130). For 3D sphere culture, hESCs were dissociated into single cells with TrypLE Select (Thermo Fisher Scientific, 12563029). ∼6×10^6^ cells in 5 ml E8 were added into one well of an ultra-low attachment 6-well plate (Thermo Fisher Scientific, 3471) and placed on an orbital shaker (within an incubator at 37°C with 5% CO_2_) at 100 rpm to induce sphere formation. The Rho-associated protein kinase (ROCK) inhibitor Y- 27632 (5 µM; Selleck Chemicals, S1049) was added to the E8 medium the first day after passaging or thawing of hESCs.

### Generation of Clonal hESC Mutant Lines

Mutant hESC lines were generated using the established iCRISPR platform and previously described methods^58^. We conducted three rounds of CRISPR, picking ∼300 clones each time, but failed to obtain biallelic (homozygous or hemizygous) S1344L mutants. Subsequently, we mutagenized PAM sites in the *TRE-Dicer1\** transgene following a previously established strategy^14^ to render it immune to gRNA targeting, so that only endogenous *Dicer1* was edited. This was transduced into H1 iCas9 hESCs^59^ and a stable clone was isolated.

Briefly, gRNA and ssDNA were added at 10 nM and 20 nM final concentration, respectively. gRNAs or gRNA/ssDNA and Lipofectamine RNAiMAX were diluted separately in Opti-MEM (Thermo Fisher Scientific, #31985070), mixed together, incubated for 5 min at room temperature, and added dropwise to the fresh iCas9 hESCs just plated. A second transfection was performed the next day to increase the gRNA transfection efficiency. Two days after the gRNA/ssDNA transfection, hESCs were dissociated into single cells and replated at a low density (700-2,000 cells per 10cm dish). The rest of cells were collected, and genomic DNA was extracted using DNeasy kit (QIAGEN, #69506). T7 endonuclease I assay was then performed to assess the CRISPR mutagenesis efficiency. ∼10 days later, individual colonies were picked, and replated into individual wells of 96-well plates pre-coated with VTN. The genomic DNA was isolated and PCR was performed for each colony to identify mutant clones.

### Growth Curves

hESCs were disaggregated using TrypLE Select and then 250000 cells were plated into one well of a 6-well plate on vitronectin in E8 medium with ROCK inhibitor. Cells were subsequently maintained in E8 with or without doxycycline for at least 4 passages. Cells were harvested after 3-4 days and counted using Vi-CELL^TM^XR cell viability analyzer (Beckman Coulter) for growth curves analysis.

### Small RNA Northern blotting

Total RNA was prepared using Trizol. 10µg RNA was run on 12%-20% urea polyacrylamide gel and transferred to GeneScreen Plus (Perkin Elmer) membrane, UV-crosslinked, and baked at 80°C for 30 minutes and hybridized with gamma-p32 labeled probes. Probe sequences are listed under oligo sequences in **Supplementary Table 7**.

### Western blotting

We assessed *Dicer1* transgene depletion and changes in Ago proteins upon removal of doxycycline using Western blotting. Antibodies used include anti-Dicer1 (1:1000 for WB, Abcam #ab14601), anti-Ago2 (1:1000 for WB, Abnova, #H00027161-M01), anti-Ago1 (1:1000 for WB, Sigma-Aldrich #MABE1121), anti-GAPDH (1:2000 for WB, Cell Signaling, #2118), anti-alpha-Tubulin (1:2500 for WB, Abcam #ab7291).

### Immunostaining

*Dicer1* mutant cells were stained as previously described^14^. Briefly, cells were fixed with 4% paraformaldehyde in PBS for ≤10 minutes, then blocked in 5% donkey serum in PBS-T for 5 minutes at room temperature for 5 minutes. Primary and secondary antibodies were diluted in blocking solution and incubated at room temperature for 1 hour. Antibodies used were anti- Sox2, Oct4, and Nanog (Abcam embryonic stem cell marker panel #ab109884).

### Small RNA library cloning

Small RNA libraries were prepared from total RNA collected from human embryonic stem cells after 7 days of doxycycline depletion. 1 µg of total RNA was used as the starting material, supplemented with 1 µL of miRNA Library QC Spike-Ins (QIAGEN) to enable absolute quantification of small RNAs. 3’ linker was adenylated in a reaction containing 200 pmol 3’ linker oligo, 4uL 10x 5’ DNA adenylation buffer, 4µL 1mM ATP, 1µL Mth RNA ligase, and water to 40µL final volume. Adenylation was performed at 65°C for 1hr and terminated at 85°C for 5 min. Precipitated, washed, and resuspended 3’ linker was used in 3’ ligation reaction containing 1µL 10x RNA ligase buffer, 2µL adenylated 3’ linker, 1µL RNase inhibitor, RNA, 1µL T4 RNA ligase 2 (1-249 K227Q), 2µL 50% PEG8000, and water to final volume of 10µL.

Ligation reactions were performed at 16°C for 4hr. The 3’ ligation products were 15% urea-PAGE purified. After elution of the RNA from gel slices with overnight incubation with shaking in 0.4M NaCl, the 3’ linker ligated RNA was precipitated and washed, and the pelleted product was used for the 5’ linker ligation by adding 0.5µL 10x RNA ligase buffer, 0.5µL ATP (10mM), 0.5µL 5’ linker oligo, 2µL 50% PEG8000, and water to 4µL final volume. The reaction was pipette mixed and incubated at 37°C for 3-5 min to re-solubilize, before incubating in a final reaction mix containing the addition of 0.5µL RNase inhibitor and 0.5 µL T4 RNA ligase 1 at 37°C for 4hr. cDNA libraries were prepared from the ligation products in RT reactions via addition of 2µL 5x RT buffer, 0.75 µL 100mM DTT, 1µL Illumina RT primer (1µM), and 0.5µL dNTPs (10mM) directly to the 5’ ligation reaction mix. The RT mix was pipette mixed and incubated at 65°C for 5 min and cooled to room temperature, placed on ice, and 0.5 µL of Superscript III RT enzyme and 0.5µL RNase OUT were added. RT reaction was carried out at 50°C for 1hr. cDNA libraries were amplified in reactions containing forward and illumine index reverse primers. Oligo sequences and reagents used for library preparation are provided in **Supplementary Table 7**.

### Small RNA library analysis

After removing adaptor sequences from the small RNA sequencing libraries using Cutadapt (v1.18), the reads were collapsed with fastx_collapser to eliminate PCR artifacts, and 5’ and 3’ end linker sequences were removed using fastx_clipper. The remaining small RNA reads were then mapped to hg38 genome assembly using bowtie (v1.3.1), and small RNA counts were normalized using spike-in reads for each library. After aligning reads to the genome, unmapped reads were subsequently mapped to the spike-in sequences and quantified for normalization. Differential expression analysis of small RNAs was conducted with the Limma package from Bioconductor in R.

### Luciferase sensor assay

The psiCHECK2 vector containing bulged or perfectly matched miRNA target sites was cloned using annealed oligos complementary to the miRNA sequence. Oligo sequences used for sensor construct cloning are listed in **Supplementary Table 7**. hESCs and HEK293T cells were transfected using Lipofectamine STEM (Thermo Fisher Scientific) and Lipofectamine 3000 (Thermo Fisher Scientific), respectively. Luciferase levels were measured using the Promega Dual-Glo Luciferase Assay Kit according to the manufacturer’s instructions. Detailed protocols for each experiment are described below:

For sensor assays monitoring 5p miRNA activity in hESCs (**Figure 2D**), the sensor and cognate targeting miRNA were co-transfected 5 days post Dox withdrawal and luciferase levels were measured 2 days later. Renilla/firefly internal control ratios were taken for both cognate miRNA and non-cognate miRNA wells, and the corresponding non-cognate/cognate ratio was taken to calculate final fold repression shown.

For sensor assays monitoring endogenous 5p and 3p miRNA activity in hESCs (**Figure 2E, 4H, 4I**), the sensor was transfected in both Dox withdrawn (5 days) and Dox-fed cells in parallel for the genotypes marked in corresponding figures. Luciferase levels were measured 2 days later and internal control renilla/firefly ratio was taken for both conditions. The ratio of these internal control values between dox fed and dox-withdrawn cells was then taken and normalized to the corresponding value of wildtype cells to calculate fold repression.

For sensor assays monitoring 5p and 3p miRNAs against cancer related sensors in Dicer1-KO HEK293T cells **(Figure 5E, 5F**), corresponding sensor and marked targeting cognate miRNA were transfected together with either WT-Dicer1 or S1344L-Dicer1 expression constructs as marked in the corresponding figures, and luciferase levels measured 2 days later. Internal control renilla/firefly luciferase ratios were taken for cognate miRNA and non-cognate miRNA wells, and the corresponding non-cognate/cognate ratio was taken to calculate fold repression.

For sample size determination, at least three biological replicates were included to demonstrate statistically significant effects. Samples were randomized and analyzed in a blinded manner.

### *In vitro* RISC assembly assay

Intact HCT116 cells or Ago2-overexpressing *Dicer1-KO* HEK293T cells were lysed in a buffer containing 100 mM potassium acetate, 30 mM HEPES-KOH (pH 7.4), 2 mM magnesium acetate, 5 mM dithiothreitol (DTT), and protease inhibitor cocktails (Roche) at 4°C. The lysates were centrifuged at 14,500 g for 15 minutes. Each sample’s lysate was then incubated with ^32^P-labeled siRNA duplexes containing a G:U wobble at the 5’ end of the sense strand, along with

0.03 μg/ml creatine kinase, 25 mM creatine phosphate, 1 mM ATP, and 10% glycerol. To stop the reactions, 2 μl of a heparin mix (60 mM potassium phosphate, 3 mM magnesium chloride, 3% PEG8000, 8% glycerol, 4 mg/ml heparin) was added and incubated for the indicated time. Notably, since robust RISC maturation was observed in Ago2-overexpressing HEK293T lysates compared to wild-type HCT116 lysates, the incubation temperature was reduced to 17 °C for the HEK293T condition to limit RISC formation, while HCT116 lysates were incubated at 25 °C. RISC assembly was analyzed using native gel electrophoresis on a 4% (40:1 acrylamide:bisacrylamide) native gel electrophoresis run at 4°C in 1x TBE at 10W.

### *In vitro* miRNA unwinding assay

For the unwinding assay, RISC assembly reactions were performed using WT or *Dicer1*lll*KO* HCT116 cell lysates as described above, except at 25°C, and quenched with 1 μl of heparin mix. Samples were then loaded onto a 15% SDS-polyacrylamide gel and subjected to electrophoresis at 4°C in 1x TBE buffer to analyze native RNA^60^.

### *In vitro* Ago2 loading assay

Duplex and nicked hairpin substrates were prepared by radiolabeling of either 5p or 3p RNA for miR-95, miR-25, miR-20a, miR-30c-2 sequences (5p RNA constituting either mature 5p RNA or 5p + loop; 3p RNA constituting mature 3p RNA). Labeled RNA substrates were loaded by 12% denaturing PAGE and purified by soaking and ethanol precipitation. Purified single-stranded substrates were then annealed overnight by heating on 95°C heatblock for 3 minutes followed by slow ramp cooling of heat block at room temperature in a styrofoam box. Final products were validated by confirmatory native PAGE (used as input control). In vitro loading reactions were performed with lysates prepared from HEK293T cells transfected with myc-Ago2 plasmid. Lysate was mixed with equal volume of 2x reaction mix, consisting of 133.4mM KCl, 16.1 mM DTT, 3.4mM ATP, 40U RNaseOUT, and radiolabeled RNA duplex. Reactions were performed at 30°C for 30 min and stopped on ice with IP/lysis buffer containing 10mM EDTA. Ago2-IP was performed using Dynabead protein G conjugated Ago2 antibody for 3 hours at 4°C with rotation. The beads were subsequently washed 3 times with lysis buffer, and resuspended in 200uL 0.5M NaCl and Ago2 bound RNA was extracted with equivalent volume of acidic phenol chloroform, and ethanol precipitated overnight at -20C. RNA was pelleted, washed with 70% ethanol, and resuspended in nuclease free water, and 6X purple loading dye (NEB) was used for loading on 15% native PAGE.

ImageJ/Fiji was used to measure total signal inside uniform size regions around bands of interest (input and product bands) and background. Background signal was subtracted from each input and product signal. The ratio of resulting product and input signal was taken, and the resulting input-normalized-values were used to calculate final ratio of nicked hairpin derived product/duplex derived product for each replicate. The average of duplex derived signal (normalized as 1) across replicates was used to plot individual duplex derived product signal.

### Ago2-eCLIP library

*Dicer1* H1 hESCs of the following genotypes (*wildtype, pm/fs, pm/pm*) were expanded under Dox-depleted conditions for 10 days. Two 15 cm dishes of confluent hESCs were used per sample. The cells were UV irradiated with 150mJ/cm^2^ UVC (254 nM wavelength) two times. The cells were then lysed on ice with iCLIP lysis buffer (50 mM Tris-HCl pH7.4, 100 mM NaCl, 1% NP-40, 0.1% SDS, 0.5% SDC, protease inhibitor cocktail (Roche), and 40 U/ml RNaseOUT (Sigma)) for 20 min on ice. TurboDNase (Invitrogen #AM2238, 10U), RNase I (0.05 U/mL) were added subsequently, and samples were incubated at 37°C for 5 min. Treated lysates were immunoprecipitated with anti-Ago2 antibodies (Abnova) or control IgG (MilliporeSigma) coupled with Dynabeads protein G (Thermo Fisher Scientific) for 2 hrs at 4°C with gentle agitation. The immunoprecipitate was washed sequentially with high-stringency buffer, high-salt buffer (50 mM Tris-HCl pH 7.4, 1 M NaCl, 1 mM EDTA, 1% Nonidet P-40, 0.5% sodium deoxycholate, and 0.1% SDS), and PNK wash buffer (20 mM Tris-HCl pH 7.4, 10 mM MgCl_2_, and 0.2% Tween-20).

The immunoprecipitated RNAs were dephosphorylated on the beads with FastAP Thermosensitive alkaline phosphatase (Thermo Fisher Scientific) and radiolabeled in 300 μl mixture: 60 μl 10X PNK buffer (NEB), 5 μl RNaseOUT, 2.0 μl γ-32P-ATP (15 μCi), 7 μl PNK enzyme (NEB M0201) and 222 μl H2O for 20 min at 37°C. The immunoprecipitated materials were washed with the high-salt buffer, and then 3’ linker ligation was carried out 2 hrs at room temperature in 25 μl of mixture: 9 μl H2O, 9 μl 50% PEG-8000 (NEB #B1004), 3 μl 10x T4 RNA ligation buffer (NEB), 0.4 μl RNaseOUT, 0.3 μl ATP (100mM), 0.8 μl DMSO, 2.5 μl T4 RNA ligase 1 (10U/μl – NEB M0204). Protein–RNA complexes were eluted from the beads, electrophoresed on 4–20% Bis-Tris NuPAGE gels (Thermo Fisher Scientific), transferred to nitrocellulose membranes (Whatman), and visualized by autoradiography.

The area of the membrane corresponding with proteins that were ∼100 kDa (the MW of Ago2) to ∼150 kDa in size was excised from each lane of the blot. Excised lanes were treated with proteinase K (MilliporeSigma) and 7M urea (MilliporeSigma). The RNA-adaptor products were purified by acid phenol:chloroform:isoamyl alcohol extraction and ethanol precipitation. Purified RNAs were then subjected to reverse transcription using Superscript III (Thermo Fisher) with adapter-specific RT primers according to the manufacturer’s instruction. Resulted single-stranded cDNA products were purified using ExoSAP-IT (Thermo Fisher Scientific), and ligated to 5’ linker in 20 μL of mixture: 1.1 μl H2O, 2 μl 10x T4 RNA ligase buffer (NEB), 1 μl of 5’-linker, 9 μl 50% PEG-8000 and 0.5 μl of T4 RNA ligase 1 (10U/μl – NEB M0204) at room temperature overnight. The linker-ligated cDNA products were purified using MyONE Silane beads (Thermo Fisher Scientific), and PCR-amplified. The resulting libraries were gel-purified and sequenced on Illumina HiSeq. Oligonucleotides used for CLIP library preparation are provided in **Supplementary Table 7**. The iCLIP library was processed using the CLIP Tool Kit^61^.

### CDF analysis of differential expression according to miRNA seed frequency

We selected the top 100 most downregulated 5p miRNAs in hotspot tumors (log2FC ≤1) (**Supplementary Table 6**) and derived a list of TargetScan defined conserved 5p sites specific to these depleted miRNAs. Each unique occurrence of a conserved site in an individual gene’s 3’UTR was counted as contributing an incremental 5p seed count of 1 to that gene’s total 5p seed count. This criterion was used to define the TEST gene sets (in increasing series of 5p seed counts) while anchoring the CONTROL set as comprising genes with no occurrence of any such 5p conserved sites. We defined test gene sets based on 3p seed occurrences by counting unique occurrences of 3p seeds across all expressed genes from the UCEC cohort, as similarly done for 5p sites, with genes containing 0 occurrence of 3p seed in their 3’UTR comprising the control set.

The following uniform analyses were performed for Dicer ES cell and TCGA UCEC data to assess contribution of frequency of 5p or 3p miRNA target sites to distribution of altered genes comparing Dicer1 hotspot cases to controls. Differential gene expression and miRNA expression analysis was performed to obtain (1) log10FC of genes as plotted along X-axis of CDF and to (2) catalogue 3’UTRs of all expressed genes according to number of occurrences of 5p and 3p target sites. Information from (2) was used to partition gene sets according to frequency of such occurrences to define control gene set (0 occurrence of miRNA target site) or test gene sets (defined number of counts as annotated in corresponding CDF plots). Cumulative distribution of control and test sets were plotted along Y-axis and Kolmogorov-Smirnov test was used to assess significance of control and test set distribution separation.

### Data availability

All of the raw sequencing data were deposited in NCBI Gene Expression Omnibus under accession GSE274849, GSE274851, and GSE274853. Original data have been provided in Source Data. All other data supporting the findings of this study are available from the corresponding author on request.

### Code availability

This paper does not report original code, as no special in-house code was required. However, any additional information needed to re-analyze the data is available upon request.

## Acknowledgments

We thank Gillian Lin and Omprakash Shriwas for participating in preliminary analysis of Dicer1 mutant hESC lines, Luke Demarco for preliminary analysis of UCEC datasets, and Arpan Das and Rui Yi for preliminary tests of Ago CLEAR-CLIP protocols. DJ was supported by CTSC-TL1 fellowship C01 5TL1 TR002386-07. SL was supported by NYSTEM contract #C32559GG and the Center for Stem Cell Biology at MSK. DY was supported by a NYSTEM training grant from the MSKCC Center for Stem Cell Biology (CSCB) (DOH01-TRAIN3-2015- 2016-00006 to D.Y.). Work in DH’s group was supported by UM1HG012654. Work in ECL’s group was supported by the Functional Genomics Initiative (FGI), the National Institutes of Health (R01-GM083300), the Tow Center for Developmental Oncology, Chan Zuckerberg Initiative (CS-2024-Lai), and MSK Core Grant P30-CA008748.

## Author contributions

DJ: biochemical and computational analyses, manuscript preparation. SL: performed biochemical and computational analyses, manuscript preparation. DY: analyzed Dicer1-mutant hESC phenotypes. RR: generated Dicer1-mutant hESCs. RS: performed biochemical analyses. DH: obtained funding, supervised hESC studies. ECL: obtained funding, supervised biochemical and computational analyses, manuscript preparation.

