## Supplementary Figures 1-10 for "Human Dicer1 hotspot mutation induces both loss and gain of miRNA function"

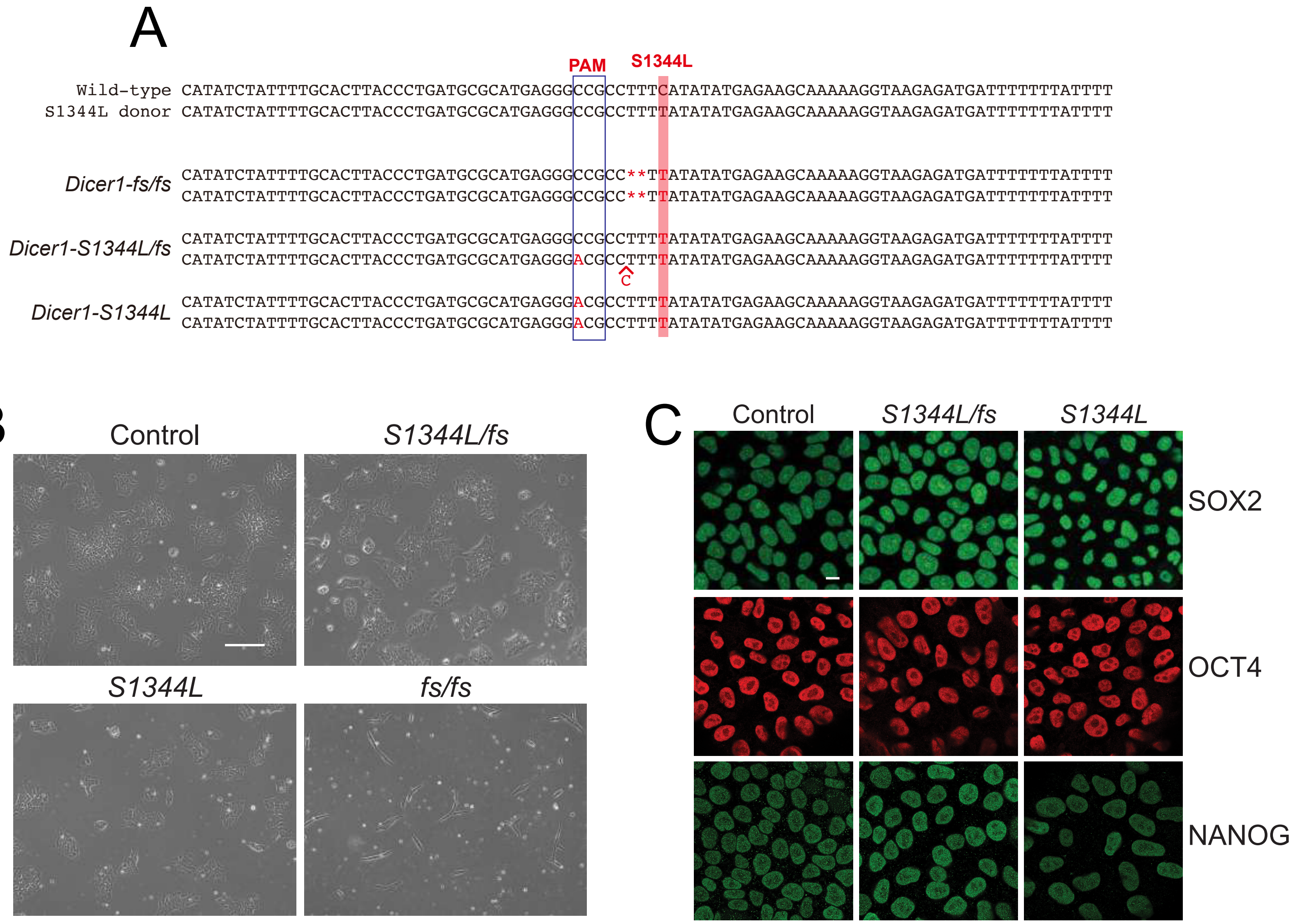


Extended Data Fig. 1

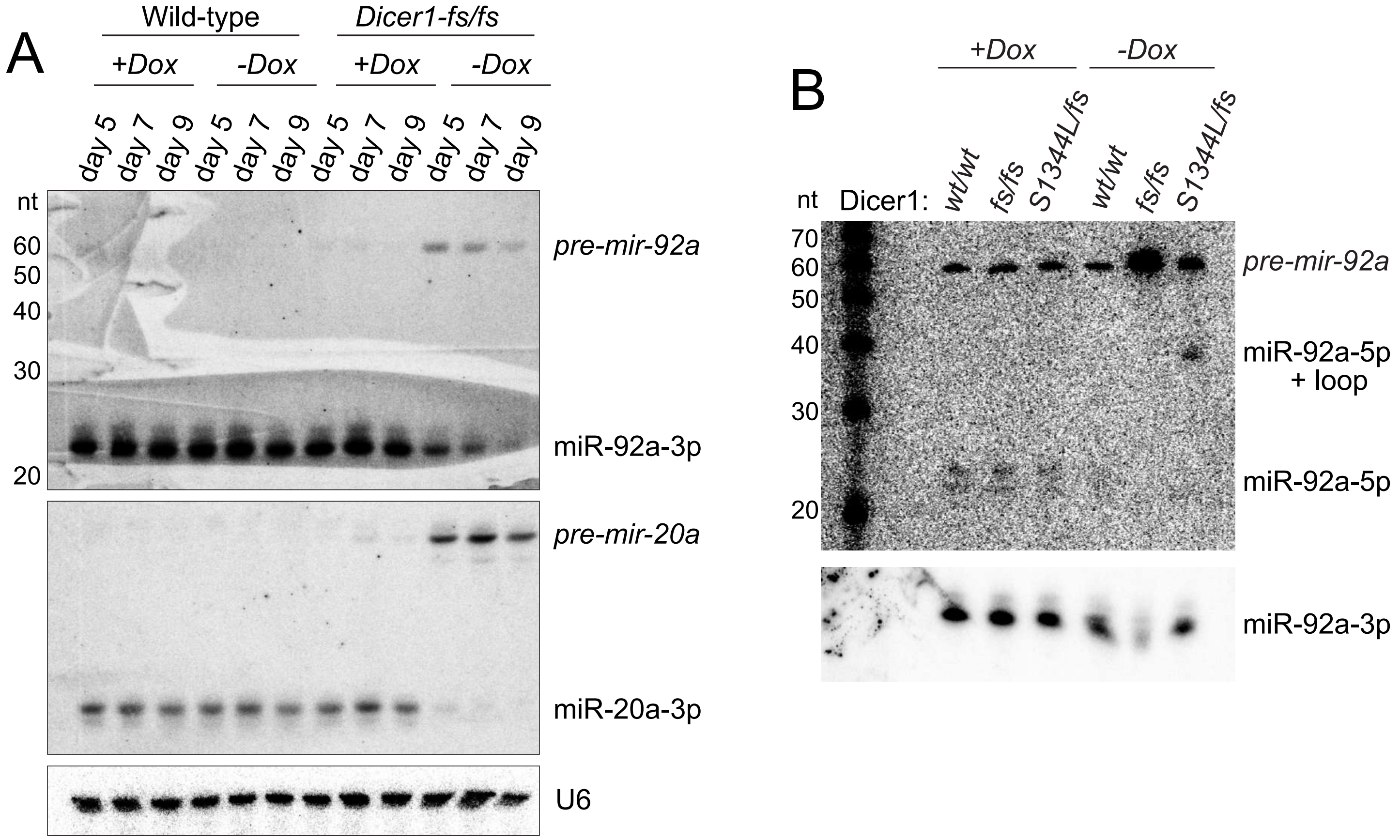


Extended Data Fig. 2

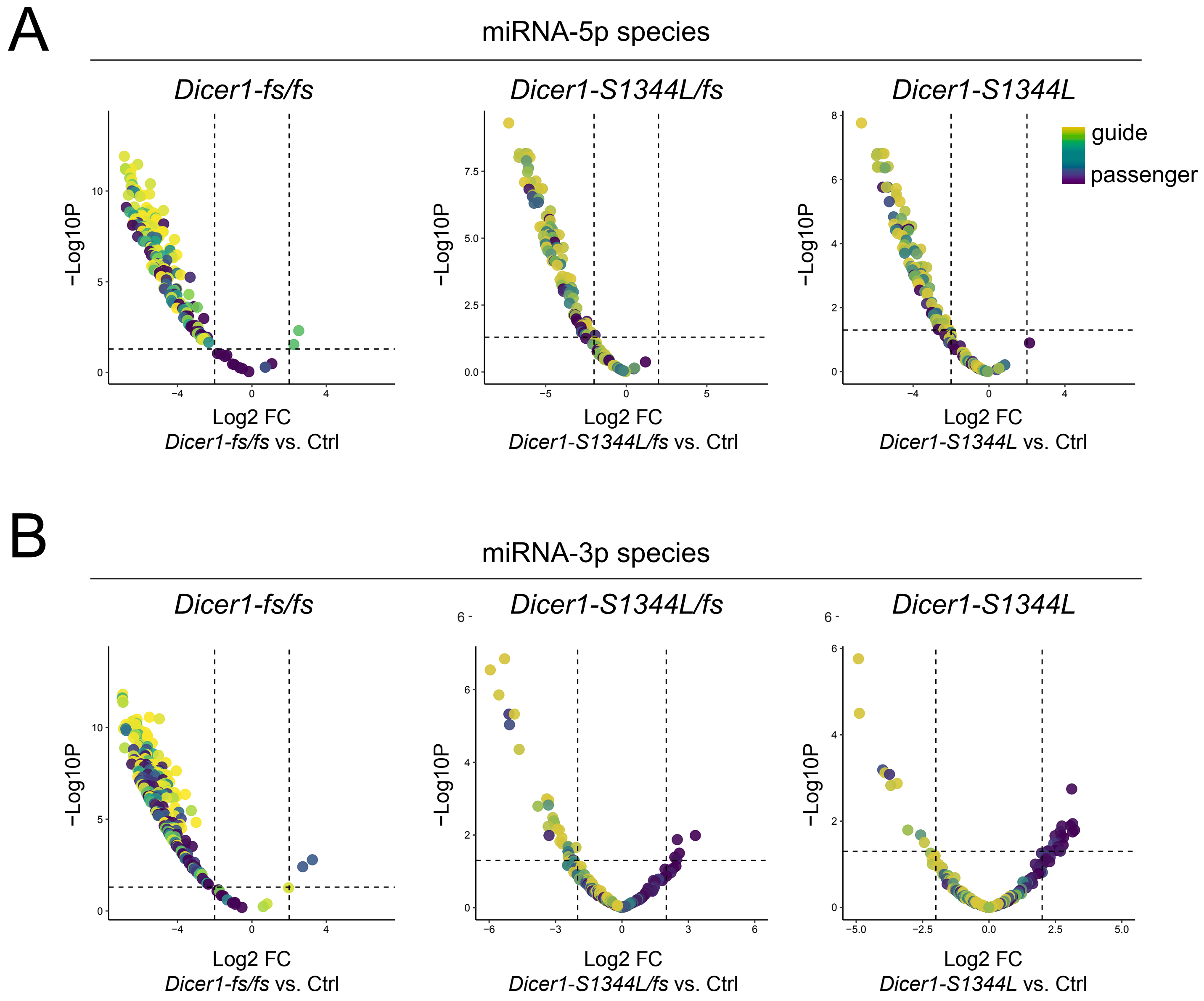


Extended Data Fig. 3

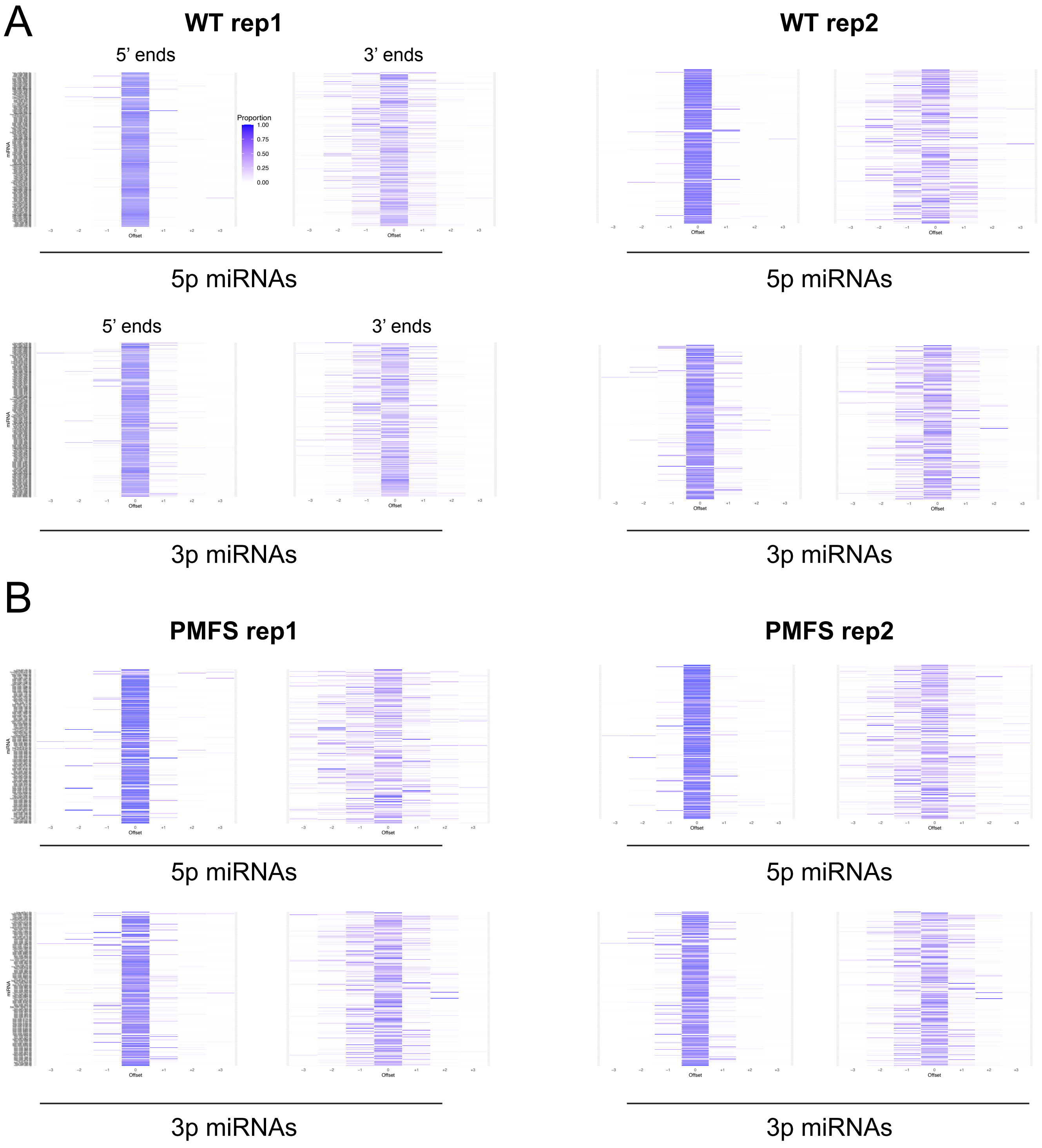


Extended Data Fig. 4

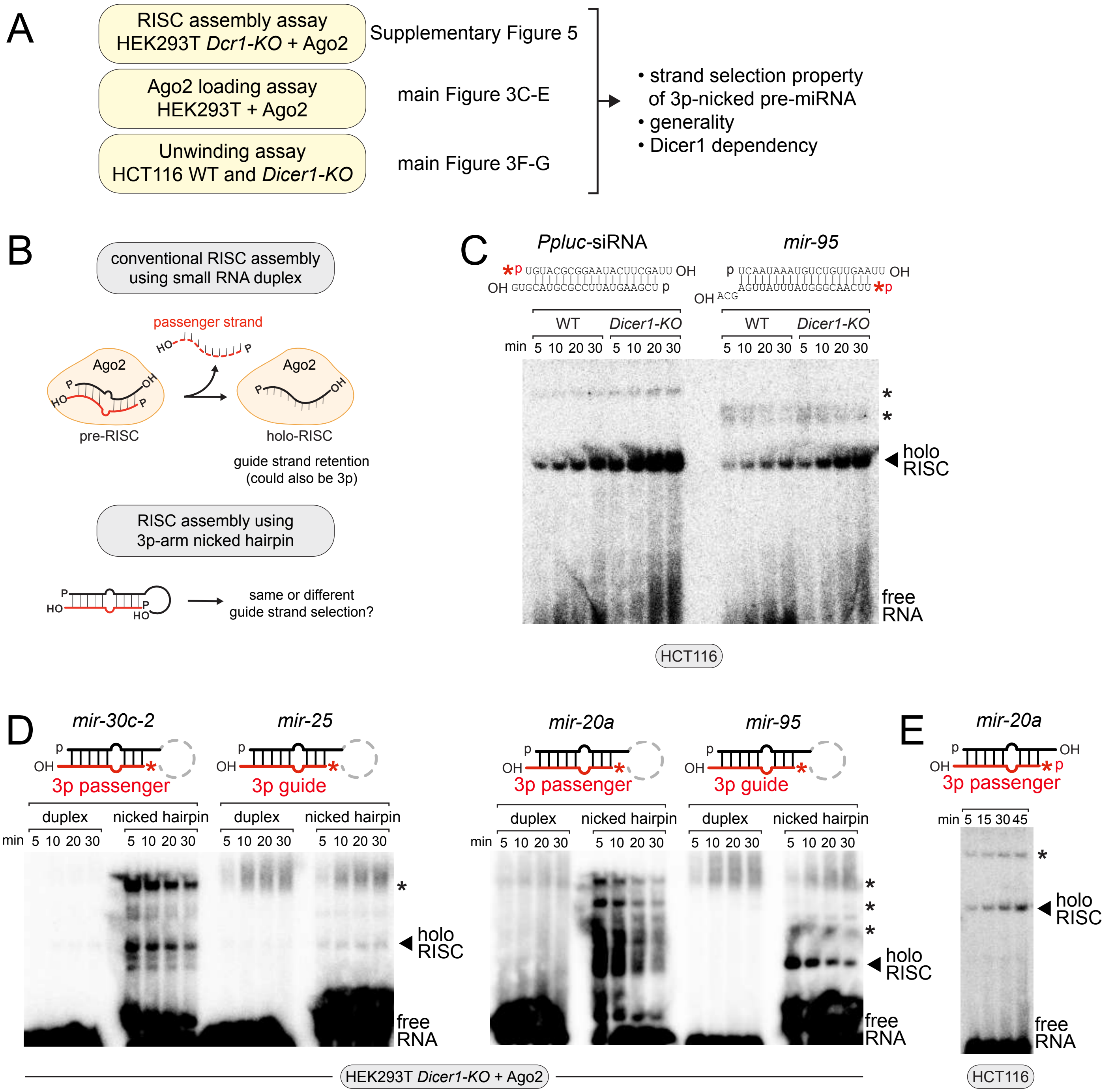


Extended Data Fig. 5

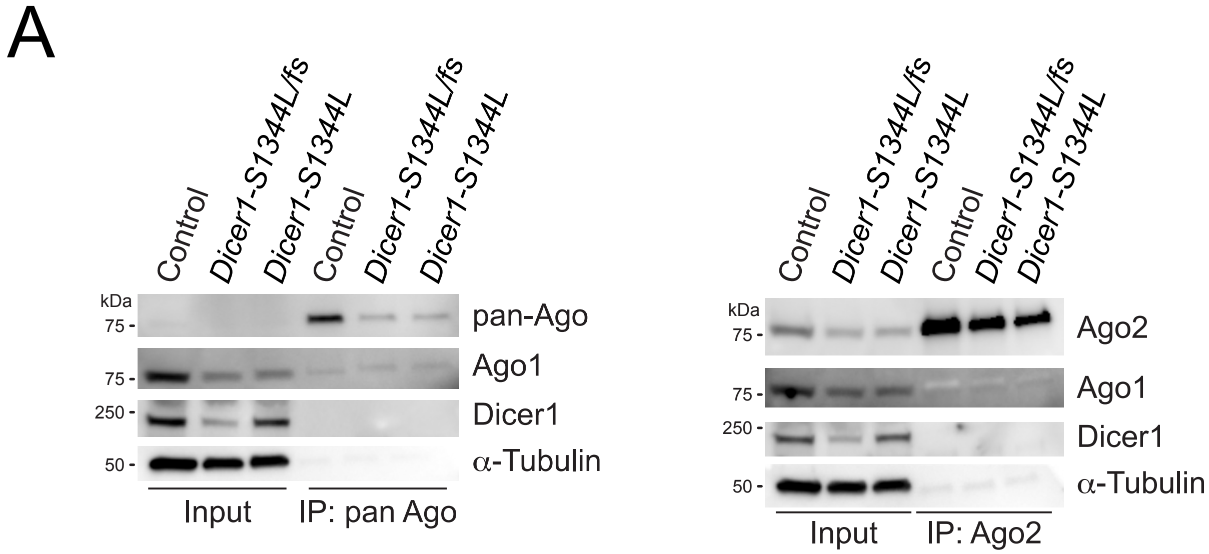


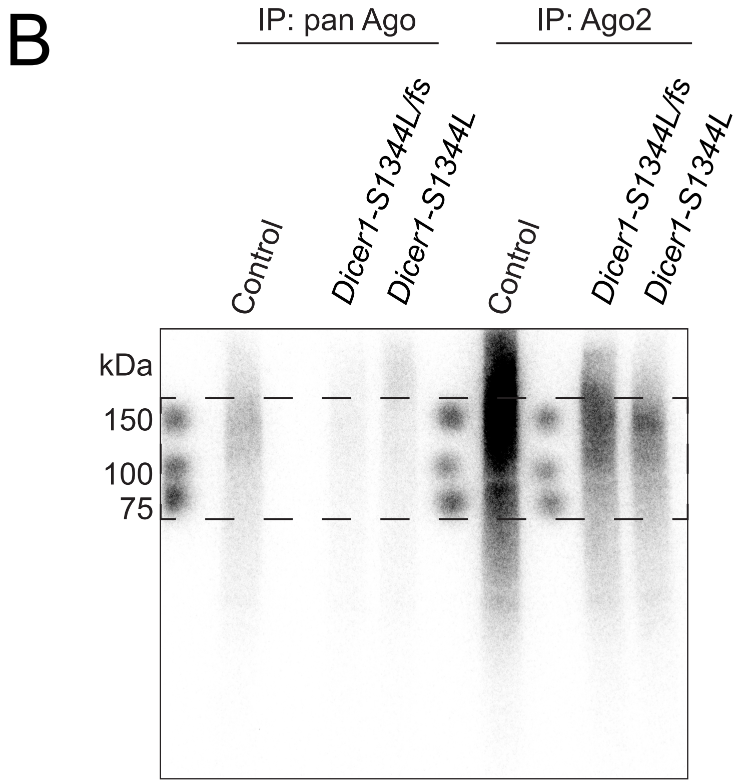


Extended Data Fig. 6

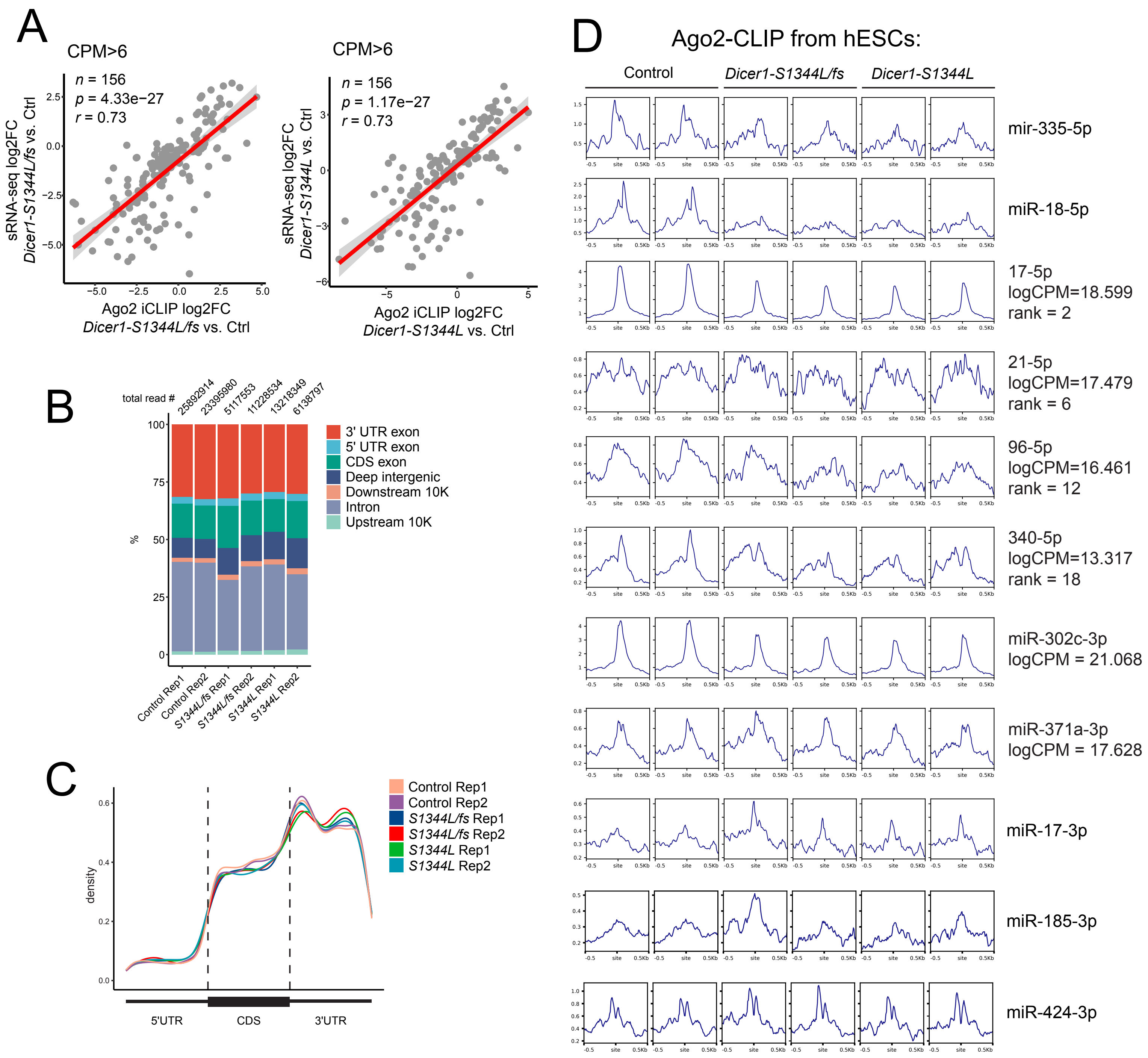


Extended Data Fig. 7

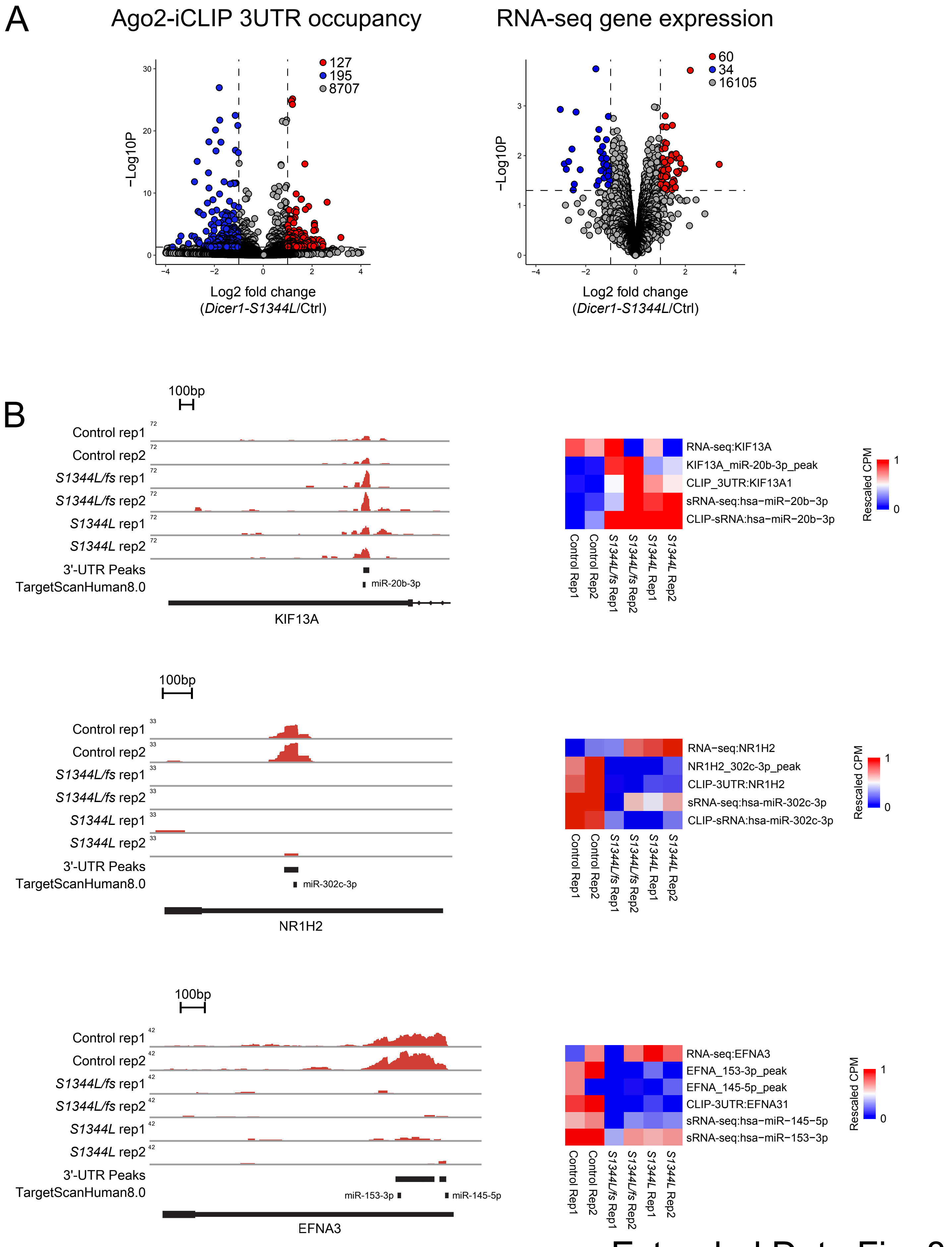


Extended Data Fig. 8

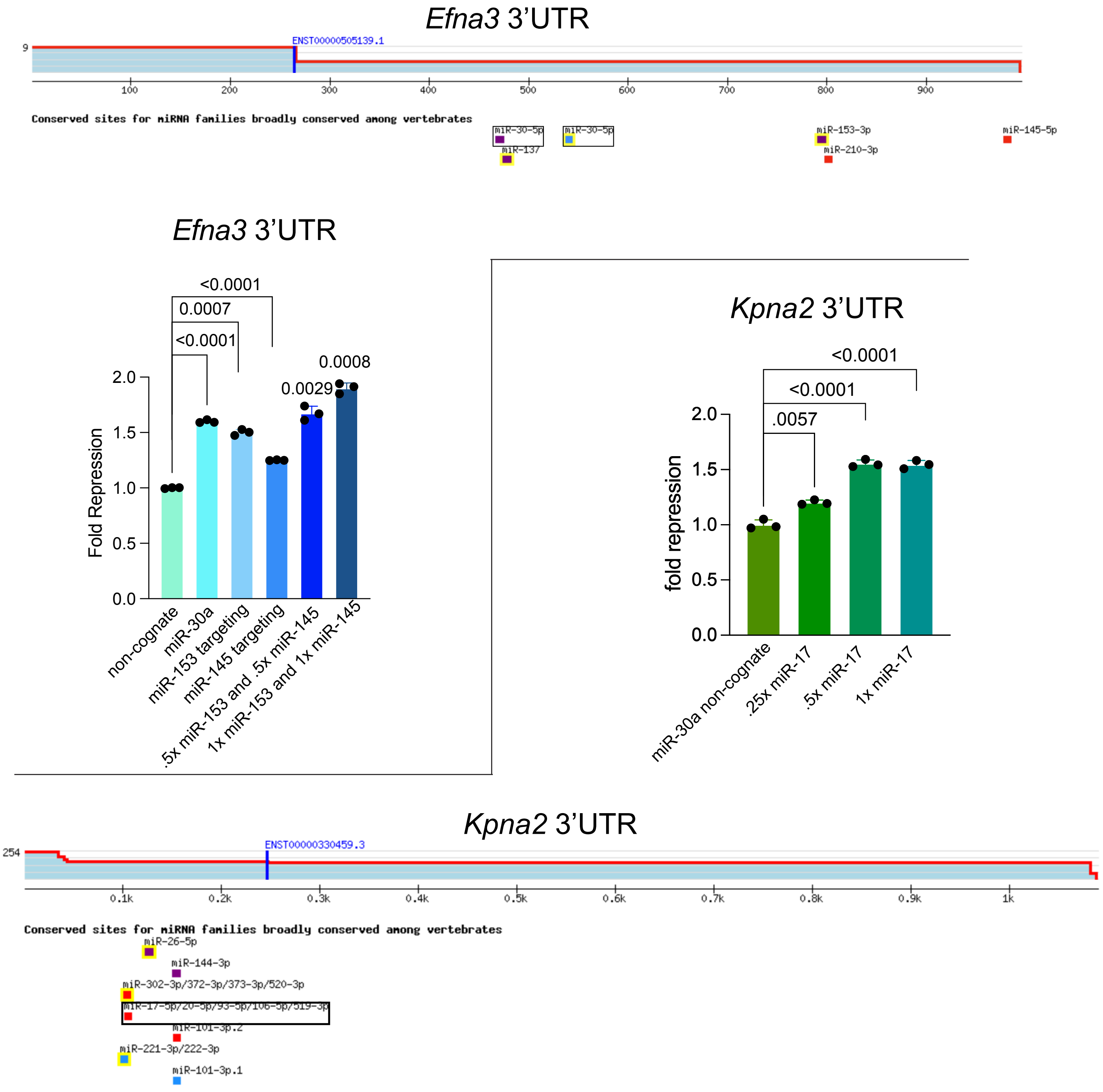


Extended Data Fig. 9

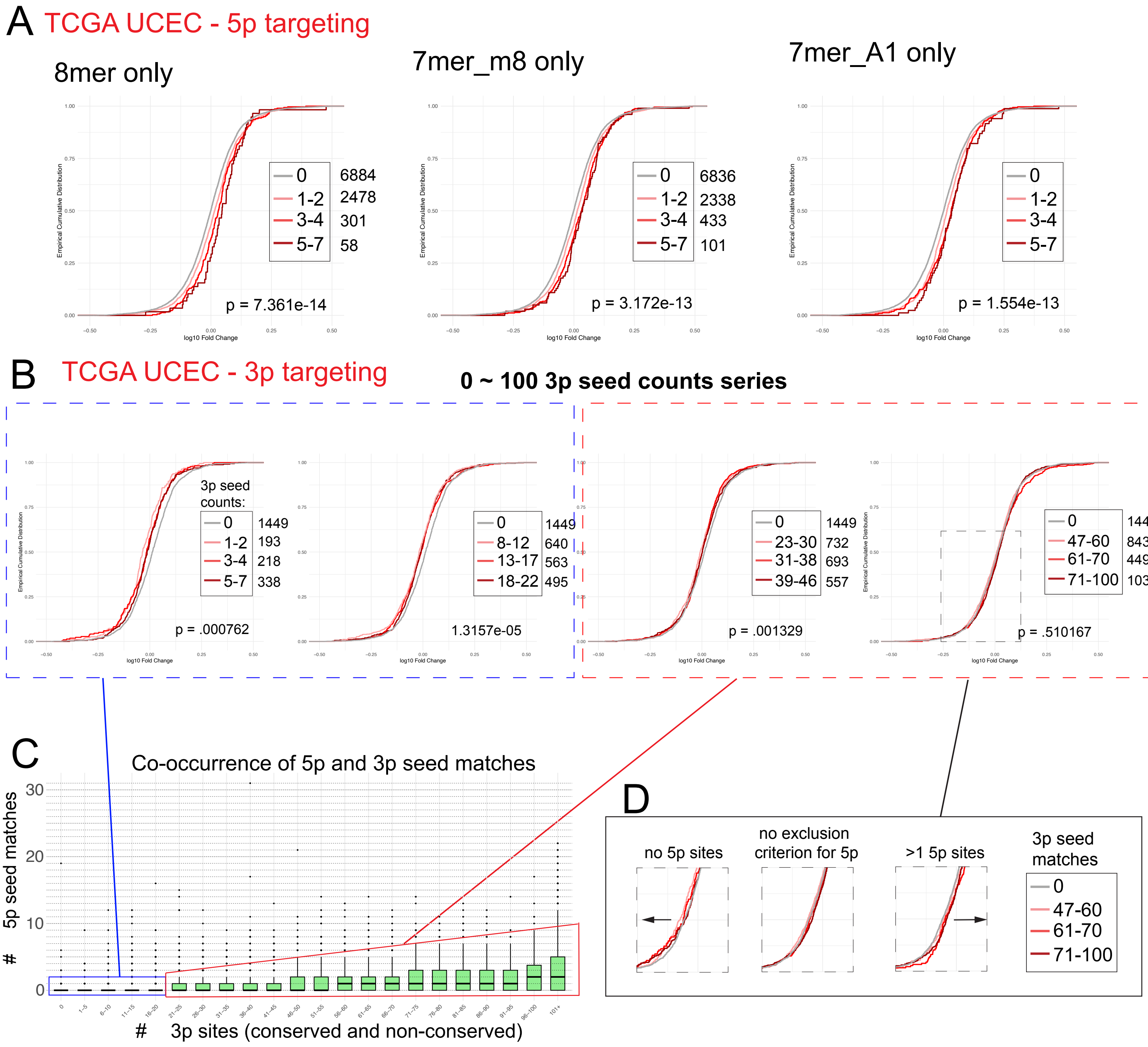


E

*CDH1* 3' UTR miR-30b-3p seed matches

ATCGGCGGCCGCGTCTCATTCTATCGGCCAGGCTGGAGTGCAGTGGTGCAATCACAGCTCACTGC
AGCCTTGTCCTCCCAGGCTCAAGCTATCCTTGCACCTCAGCCTCCCAAGTAGCTGGGACCACAGG
CATGCACCACTACGCATGACTAATTTTTTAAATATTTGAGACGGGGTCTCCCTGTGTTACCCAGGCT
GGTCTCAAACTCCTGGGCTCAAGTGATCCTCCCATCTTGGCCTCCCAGAGTCTCGAGCGAT

*PLA2R1* 3' UTR miR-17-3p seed matches

AAATATTTCTTCTCTTGAAGGGTCCACTCCTTATTAAAGATAAGATTGAACTGAATGTTTACTTAAACG
TTCAGTCTTTTGGAGTTTCTTTACTCTTCTCAAAATGGTGGTTGGATATATAAGAACTTGCTGTGTCC
TGGTAGTGTTGAGGGGAGAAGGTGACAACTTACTGTGGATGTATGTTAGCTGGTGTCCACTGAGAA
TAGCTGTCTGGCCTCACTCCTGGGAATTCTGCTGGCATCTGCTGTGAAGTCTGCAGTTGCTTTAAT
CTGTAGTCTGTTCTCTTTGATTTGGCCTCATTCTGGCTGTCCCTGGTGCTGTCTTATCTCTTGAGAA
CTGGCTGTGTCTCCTGTTGAATTCCCTATCCGAACACTGCAGTTCTTGGCTACAAGGCAGTCACTT
CACCCCTGCAGTGGTGAGATCCCATTAGCTACCATTAGCTACTCTTTTCGTCCAGAAAGGCGAAAT
GCCATAGCAAACAATCCCACAGTCTTGGTGGATTAACACAGCAAAGGTTTATTTCTCACCCACACCT
GCATGGGCTGGCAAGAGGACTTTGCTCTGTTCAGTCATTCATGGGTCCATGCTTCTCCTATATAGCC
AGCCCTCTGTCTCCTAAGGCCTGGAGTCCTCCATGGGCCTCCAATCTGTGG

*CIITA* 3' UTR miR-30b-3p seed matches

TCAACCTCCTGGGCTAAGTGATCCTCCCACCTCAGCCTCCCGAATAGCTGGGACTACAGGTGTGAG
TCACCAAGCCCAGTTAATCTTTAGTTTTATTTTTGTAGAGCCAGGGTCTCACTATGTTGCCCAGGCAG
GTCTTGAACTCCTGGCCTCAAGTGATTCTCCTGCCTCAGCCTCCCAAAGTGCTGGGATTACAGGTG
TGAACCACCACACCCAGCCCACTTCTGCCATATTCTGTTGGCCAGTGTGACAAGGATTGCTACTGT
CCTACCCACCCTCCTTTCACCACATGTGCACATGCACGTGTGTGCACGTACACACACATACACACA
CACGCGTGCACACACCAGAGCCCACCTTGGCTCAAGTCCTCTTTTCTGAGAGGACTTTTCTTTGTG
GCTTCCTAAAATTCAGTGGAAAATTAATTGTTTGGGGATGAGAAAGGTTGAAGCACCAAAAGCTTAC
CAAGGGGAATGTTTGCCTCTGCCTCTGACACACAGCTCTGTCCTGGGAGTGAGCTGGTTTTTAGCG
GAGACGGAGTCCCACCTTGGCTGCAGGGAGTCCGAAAGGGACTTGGAAGCTCCGCTTTCTACCCA
GCGAGCTGCCGGCACGACTGTTGCTTTTCACTCGGGCATAGTCTGCTCAGAAGCCCCAATTCAAG
ACATCTTAGCTTACTCCTGGTGGCAGTGGGAGCTGCCTGTTCAGCTCCAGCTCACCAGCCCCAGT
GCCCACAGGATCAGTCTGATTCCCAAGCTCTGCTCCTCTCCCCAGCAAGTGAGAGCTGGGTGTCA
AGAGGGTCTGAGGAATCCAGACCCGGCTGCGTGGTTGATGATCTTTGCAGACAGGCAAGCACCTC
CTGCCTCGAACTGGCTGCAAAGGGTATCAGGTGCTGCCATCCAGGTTCAGCTTGTAAATAGCTCAG
GTGTCACCCTGCAAGGGATCCCTGCACTTGCAGACTCTACAGAGGCCATGGGCCTCCTGTGTGTG
CGCCTACAGGAGTGAGCACTAAGGTGTGCTCTGATCATCCACTGTGCCATGTGCCAGGTTTTTGGT
CTGACCCACTGCTTTTGGTCTGTGTGATGATGAAATGAGGTCAGAGCCTGCAGTTATTAACACTAAC
GATTTGTTAATGATAGCTACTGTTTATTATTTTATTTAATTTTTTTGAGACAGAGTTTTCACTCTTATTGC
CCACGCTGGAGTGCAGTGGCATGATCTCGGCTCACTGCAACCTCTGCCTCCTGGGTTCAAGCAAT
TCTCCCACCTCAGCCTCCCAAGTAGCTGGGATTACAGGCATGCACCACCACGCCTGGCTAATTTTT
GTATTTTTAGTAGAGATGGAGTTTCACCATGTTGGCCAGGCTGGTAATCTGCCTGCCTTGGCCTCCC
AAAGTGCTGGGATTACAGGGGTGAGCCACCGCACCCGGCCACTAC

Extended Data Fig. 10
